# HLA-A*11:01-Restricted T Cell Receptors Targeting Mutant NPM1 for Adoptive Cell Therapy in Acute Myeloid Leukemia

**DOI:** 10.64898/2026.09.25.754423

**Authors:** Aidan J. Pursley, Uns Kirmani, Amanda L. Chen, Zhiya Yu, Kenichi Hanada, Catherine M. Ade, Christopher D. Chien, Nuri Cha, Willy A. Flegel, Peter D. Aplan, Martin Carroll, Naomi N. Taylor, James C. Yang, Steven Z. Pavletic, Sung-Yun Pai, Vid Leko

## Abstract

Acute myeloid leukemia (AML) is an aggressive hematologic malignancy associated with a poor prognosis and an unmet need for more effective treatment options, particularly after relapse. One potential approach to address this need, which has been used successfully in solid tumors, is adoptive transfer of T cells engineered to express specific T cell receptors (TCRs). Particularly attractive are TCRs that can target epitopes derived from oncogenes on which cancer cells depend while sparing normal cells that do not harbor the mutation. One promising oncogenic target in AML is mutated nucleophosmin 1 (mtNPM1), present in up to 30% of AML cases. Most NPM1 mutations result in a common frameshifted sequence that generates peptides presented by several class I human leukocyte antigens (HLAs), such as HLA-A*11:01. In this study, we vaccinated HLA-A*11:01-transgenic mice with the mtNPM1 peptide AVEEVSLRK and isolated six functional TCRs from mtNPM1-reactive lymphocytes. When engineered into human T cells, two of these TCRs consistently mediated robust and specific lysis of an AML cell line and primary AML cells co-expressing HLA-A*11:01 and mtNPM1. Neither TCR recognized related human peptides, HLA-A*11:01-expressing normal peripheral blood mononuclear cells or stem/progenitor cells, or cells expressing diverse allogeneic HLAs, suggesting a low risk of cross-reactivity and alloreactivity. In an AML xenograft model, both TCRs controlled AML in hematopoietic organs, although extramedullary tumors emerged during treatment, potentially reflecting the aggressive behavior of the model that may not be representative of AML in patients. Together, these data suggest that the two TCRs represent strong candidates for further preclinical development in adoptive T cell therapy for NPM1-mutated, HLA-A*11:01-positive AML. These results also support the use of HLA-transgenic mice as a platform for generating TCRs targeting oncogene-derived epitopes in hematologic malignancies.

## Introduction

Acute myeloid leukemia (AML) is an aggressive hematologic malignancy characterized by the clonal expansion of immature myeloid cells in the bone marrow, peripheral blood, and other tissues. Despite advances in therapy, the overall 5-year survival rate remains approximately 33%^1^. For newly diagnosed AML, intensive chemotherapy and allogeneic hematopoietic stem cell transplantation (allo-HSCT) can be curative but are associated with substantial treatment-related morbidity and mortality and are not suitable for many patients. For patients who relapse, fewer effective options exist and outcomes remain poor^2^. Thus, more effective and less toxic therapies are needed for patients with AML.

One potential approach for the treatment of this disease is to engineer tumor-reactive T-cell receptors (TCRs) into autologous T cells from peripheral blood for infusion back into the patient. This form of adoptive cell therapy, known as TCR-engineered T-cell therapy (TCR-T), has induced objective responses in patients with treatment-refractory solid tumors by targeting both tumor-associated antigens (TAAs) and neoantigens^3–7^. In AML, TCR-T targeting the TAA Wilms’ tumor 1 (WT1) has demonstrated encouraging preliminary clinical activity in preventing relapse after allo-HSCT^8^. However, the clinical efficacy of targeting neoantigens with TCR-T in AML has not yet been reported.

Unlike TAAs, which may also be expressed in normal tissues and therefore confer a risk of on-target, off-tumor toxicity, neoantigens arise from tumor-specific genomic alterations that are absent from normal tissues, allowing more selective targeting. Neoantigens derived from mutated oncogenes are particularly attractive therapeutic targets because they are generally recurrent (i.e., shared among patients), clonally expressed, and, by contributing to maintenance of the malignant phenotype, less likely to be lost from the tumor under therapeutic pressure^9^.

Mutations in nucleophosmin 1 (NPM1), which occur in up to one-third of AML cases, generate one such target. The vast majority of NPM1 mutations consist of four-base-pair insertions in the terminal coding exon, resulting in a frameshifted mutant protein that aberrantly localizes to the cytoplasm (from which its alternative designation, NPM1c, is derived) and contributes to maintenance of the leukemic state^10^. The most common NPM1 mutation is type A, characterized by a TCTG duplication and accounting for approximately 70–80% of mutant cases^11^. This and several rarer mutation types (including D, G, and H) generate the shared C-terminal sequence CLAVEEVSLRK, which generates peptides naturally presented by several class I human leukocyte antigen (HLA) molecules (Figure 1A)^12,13^. The C-termini of additional mutation types (B and C) have distinct upstream sequences but also contain the shared AVEEVSLRK sequence.

**Figure 1.**
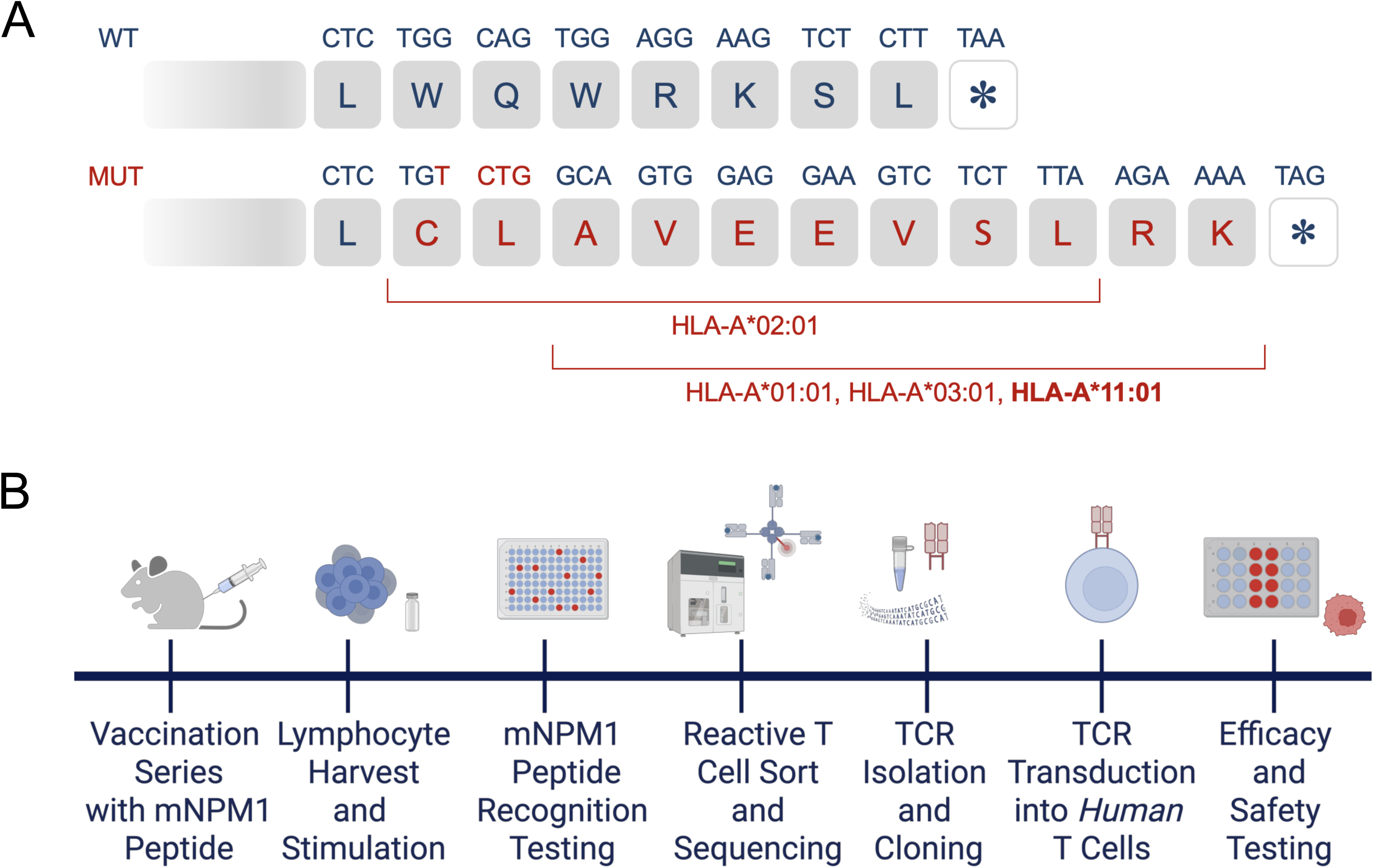
NPM1 mutation-derived epitopes and workflow for TCR identification using HLA-A*11:01-transgenic mice. **(A)** Recurrent four-base-pair insertions in exon 12 of NPM1 affect up to 30% of AML cases. These include the common type A (TCTG) insertion, which generates a shared frameshifted C-terminal sequence. This sequence contains the neoepitope AVEEVSLRK, which binds to HLA-A*11:01 and serves as the target for the TCRs described in this study. This peptide was also shown to bind to HLA-A*01:01 and HLA-A*03:01. The CLAVEEVSL peptide, which binds to HLA-A*02:01 and is targeted by a TCR currently undergoing clinical evaluation, is shown for reference. **(B)** Overview of the vaccination approach using HLA-A*11:01-transgenic mice and the AVEEVSLRK peptide to isolate reactive TCRs. The image was created with BioRender.

The mutant NPM1 protein generated by these mutation types, referred to here as mtNPM1, therefore represents an attractive target for immunotherapy. An HLA-A*02:01-restricted TCR targeting the CLAVEEVSL epitope is currently being evaluated in a phase I/II clinical trial for patients with relapsed or refractory AML (NCT06424340). However, clinically useful TCRs targeting AVEEVSLRK remain limited. A TCR recognizing this peptide presented by HLA-A*11:01 has previously been identified but demonstrated insufficient antileukemic activity in a preclinical model, suggesting that additional receptors are needed^12^.

Many TCRs considered for clinical development across different cancers have been isolated from patients or HLA-matched healthy donors using *in vitro* sensitization methods^12,14^. Alternatively, transgenic mice that express HLA proteins provide a robust platform for generating high-avidity TCRs, several of which have been used in clinical trials for solid tumors^15–17^.

In this study, we sought to identify mtNPM1-reactive TCRs for the treatment of HLA-A*11:01-positive AML. To do this, we vaccinated HLA-A*11:01-transgenic mice with the AVEEVSLRK peptide and isolated their lymphocytes for downstream TCR sequencing and functional testing (Figure 1B). We identified two TCRs that specifically recognized and lysed AML cells co-expressing mtNPM1 and HLA-A*11:01, controlled leukemia in the hematopoietic compartments of AML-engrafted mice, and demonstrated favorable preclinical safety profiles.

## Methods

### Vaccination of HLA Transgenic Mice

HLA-A*11:01-transgenic mice on a C57BL/6 background were procured from Taconic Biosciences (Germantown, NY). Four 4-week-old female mice each received a total of three vaccinations with mtNPM1 peptide emulsions. To prepare them, 1 mg of mtNPM1 peptide (AVEEVSLRK; GenScript, Piscataway, NJ) and 1.2 mg of hepatitis B virus core antigen helper peptide (128–140, TPPAYRPPNAPIL; GenScript), both supplied as lyophilized powders, were each reconstituted in 50 μl DMSO (Sigma-Aldrich, St. Louis, MO). The two peptide solutions were then combined and diluted with 400 μl PBS (Gibco, Waltham, MA) to a final volume of 500 μl.

For the first vaccination, peptide solution was emulsified in 500 μl Complete Freund’s adjuvant (Sigma-Aldrich, St. Louis, MO) using a Dremel 4200 rotary tool equipped with a smooth, straight stainless-steel probe (Dremel, Mount Prospect, IL), and 50 μl of the emulsion was administered into each footpad. Subsequent peptide solutions were emulsified in 500 μl Incomplete Freund’s Adjuvant (Sigma-Aldrich, St. Louis, MO), and 100 μl per mouse was administered subcutaneously in the base of the tail. Intervals between individual vaccinations were two and three weeks, respectively. One week following the last vaccination, spleens and lymph nodes were harvested from the mice for processing and *in vitro* stimulation. All mouse studies were approved by the NCI Animal Care and Use Committee.

### *In Vitro* Stimulation of Murine Lymphocytes

Following tissue harvest, single-cell suspensions were prepared by mechanically dissociating spleens and lymph nodes through 40-µm strainers (Corning Incorporated, Corning, NY) and rinsing with 5 mL ACK lysis buffer (Gibco, Waltham, MA) or 5 mL PBS, respectively. Next, cell suspensions were washed with PBS and resuspended in complete mouse T cell medium, which consisted of RPMI 1640 medium supplemented with 10% fetal bovine serum (Cytiva, Marlborough, MA) as well as 25 mM HEPES, 1% antibiotic-antimycotic (Anti-Anti), 1 mM sodium pyruvate, 1% MEM-nonessential amino acids, 1% GlutaMAX, 55 µM 2-mercaptoethanol (all from Gibco), and 30 IU/mL recombinant human IL-2 (rhIL-2, Prometheus Laboratories, San Diego, CA). Cells were then plated in 24-well plates at a density of 2 × 10^6^ cells/well, and stimulated *in vitro* by adding the synthetic peptide AVEEVSLRK directly to the cell suspensions at 10, 100, or 1,000 ng/mL, with eight wells receiving each concentration. Cultures were then incubated for 10 days at 37 °C; a half-medium change was performed on day 5 of stimulation.

### Identification of AVEEVSLRK-Reactive Murine T Cells

To serve as antigen-presenting cells, human leukemia cell line K562 (ATCC, Manassas, VA) was first transduced to express HLA-A*11:01 as previously described^15^. These cells, hereafter referred to as K562-A*11:01 cells, were then pulsed with the synthetic peptide AVEEVSLRK at 100 or 1,000 ng/mL, or with DMSO (control) for 4 hours. Cells were then washed and plated in 96-well plates at 5 × 10^4^ cells/well in complete mouse T cell medium. Murine lymphocytes that had completed 10 days of *in vitro* stimulation were washed to remove IL-2 and added to these plates at 1.5 × 10^5^ cells/well. Co-cultures were incubated overnight at 37 °C and 5% CO_2_. The following day, murine interferon gamma (IFN-γ) production was measured as described in the following section.

### Assessment of IFN-γ Production from Murine and Human T Cells

IFN-γ production was measured from murine T cells using mouse IFN-γ ELISA. For this, co-culture plates were first spun at 400 × *g* for 3 minutes at room temperature; supernatants were then assayed using a mouse IFN-γ ELISA kit (Duoset, R&D, Minneapolis, MN), following the manufacturer’s instructions. The plates were read using the Infinite F50 Plus spectrophotometer and analyzed using the accompanying Magellan data analysis software (both from Tecan, Männedorf, Switzerland).

In experiments with TCR-transduced human T cells, generated as described below, IFN-γ production was assessed by either human IFN-γ ELISpot or U-PLEX, an electrochemiluminescence-based assay. For IFN-γ ELISpot, co-cultures were prepared in multiscreen 96-well plates with hydrophobic PVDF membranes (MilliporeSigma, Burlington, MA), pre-coated with an anti-human IFN-γ capture antibody (clone 1-D1K; Mabtech, Nacka Strand, Sweden). The next day, cell suspensions were removed and the plates were processed using biotinylated anti-human IFN-γ detection antibody (clone 7-B6-1; Mabtech) and streptavidin-ALP (Mabtech). They were developed using KPL BCIP/NBT substrate solution (SeraCare Life Sciences, Milford, MA). The reaction was stopped by rinsing with cold tap water; plates were then scanned and counted using an ImmunoSpot plate reader (Cellular Technologies Limited, Shaker Heights, OH).

For human IFN-γ U-PLEX assays, co-culture plates were spun at 400 × *g* for 3 minutes at room temperature. Cell pellets were resuspended in PBS supplemented with 0.5% FBS and stored at 4 °C for subsequent flow cytometric analysis. Supernatants were analyzed using a human IFN-γ U-PLEX kit (Meso Scale Diagnostics, Rockville, MD) according to the manufacturer’s instructions. Plates were read on a MESO Sector S600 reader (Meso Scale Diagnostics) and analyzed using the accompanying software.

### Flow Cytometry

For all experiments, unless otherwise specified, cells were stained with antibodies diluted at a 1:100 ratio in PBS with 0.5% FBS for 30 minutes at 4 °C; they were washed with PBS with 0.5% FBS before flow cytometry analysis. Antibodies were acquired from BD Biosciences (La Jolla, CA) and BioLegend (San Diego, CA) and included anti-mouse CD3 (clone 17A2), CD8 (53-6.7), and TCRβ (H57-597), as well as anti-human CD3 (SK7 or UCHT1), CD8 (SK1), CD4 (SK3), CD137 (4-1BB; clone 4B4-1), CD34 (581), CD19 (HIB19), CD14 (M5E2), CD16 (3G8), CD56 (NCAM16.2), and CD45 (HI30). Flow cytometric acquisition was done using a NovoCyte Flow Cytometer (Agilent Technologies, Santa Clara, CA), while cell sorting was performed on an SH800S Cell Sorter (Sony Biotechnology, San Jose, CA). For live-vs-dead discrimination, 7-AAD and DAPI (4′,6-diamidino-2-phenylindole), both from BD Biosciences, were used on the NovoCyte and SH800S instruments, respectively. Data were analyzed using FlowJo (Tree Star, Ashland, OR).

### Isolation of mtNPM1-Reactive Murine T Cells

To isolate mtNPM1-reactive cells from cultures that produced IFN-γ in response to AVEEVSLRK, leftover cells were first stained with HLA-A*11:01-mtNPM1 tetramers, followed by single-cell sorting of tetramer-positive populations.

To prepare HLA-A*11:01 tetramers, 45 µg AVEEVSLRK peptide and 10 µg of Flex-T HLA-A*11:01 Ultraviolet Exchange monomer (BioLegend, San Diego, CA) were combined and incubated on ice for 2 hours under UV light (366 nm wavelength). Following peptide exchange, 0.8 µg of streptavidin-PE or streptavidin-APC (BioLegend, San Diego, CA) was added four times, every 15 minutes for 1 hour. PBS was used to bring tetramer volumes to 100 µL.

For single-cell sorting, reactive murine T cells were stained with tetramers diluted 1:20 in PBS containing 0.5% FBS for 30 minutes at 4 °C. Cells were then washed to remove unbound tetramer and stained with anti-mouse CD3 and CD8 antibodies. Live, singlet, CD3^+^, CD8^+^, tetramer^+^ cells were sorted into individual wells of a 96-well PCR plate using the SH800S Cell Sorter for single-cell TCR sequencing.

### Single-Cell TCR Sequencing and Cloning of Murine TCR

Single-cell TCR sequencing and cloning were performed as previously described^15^. Briefly, sorted single T cells were subjected to cell lysis, first-strand synthesis, template switching, and cDNA preamplification using a modified Smart-seq2 method^18^. Partial TCRα and TCRβ sequences were then amplified from cDNA using the template-switch oligo primer together with primers specific for the TCRα and TCRβ constant regions (5′-GGCCCCATTGCTCTTGGAATC-3′ and 5′-GTCCGTGCTGACCCCACTGT-3′, respectively). Amplified α and β products were submitted for Sanger sequencing using 5′-CTGTCCTGAGACCGAGGATC-3′ and 5′-GTTTGTTTGCAATCTCTGCTTTTGATG-3′ primers, respectively (Psomagen, Rockville, MD). Candidate TCRs for full-length cloning were selected by mapping partial TCRα and TCRβ sequences back to their wells of origin and selecting the α/β pairs (i.e., clonotypes) identified in more than two cells.

For TCR cloning, cDNA from selected wells underwent full-length TCRα and TCRβ chain amplification by PCR using primers specific for the individual α/β chains. Amplified chains were then individually cloned into the MSGV1 vector using the In-Fusion Cloning Kit (Takara Bio, Kusatsu, Japan), according to the manufacturer’s instructions. Following cloning, individual colonies were picked and used for plasmid isolation. Plasmids were verified by Sanger sequencing, and those containing the correct sequences were pooled and used for the screening experiments.

### Retroviral Transduction of Murine TCRs into Healthy Donor PBMCs

For the screening experiments shown in Figure 3 and Supplementary Figure 2, T cells were transduced with separate TCRα and TCRβ retroviral supernatants corresponding to the same receptor. For subsequent experiments, to simplify TCR production, the α and β chains of TCR6 and TCR7 were cloned together into the same MSGV1 vector using a P2A ribosomal skip element to separate the chains (GenScript). In selected experiments, plasmids encoding HLA-A*01:01- or HLA-B*40:01-restricted TCR α/β chains recognizing epitopes derived from unrelated oncogenes were used interchangeably as irrelevant TCR controls.

**Figure 2.**
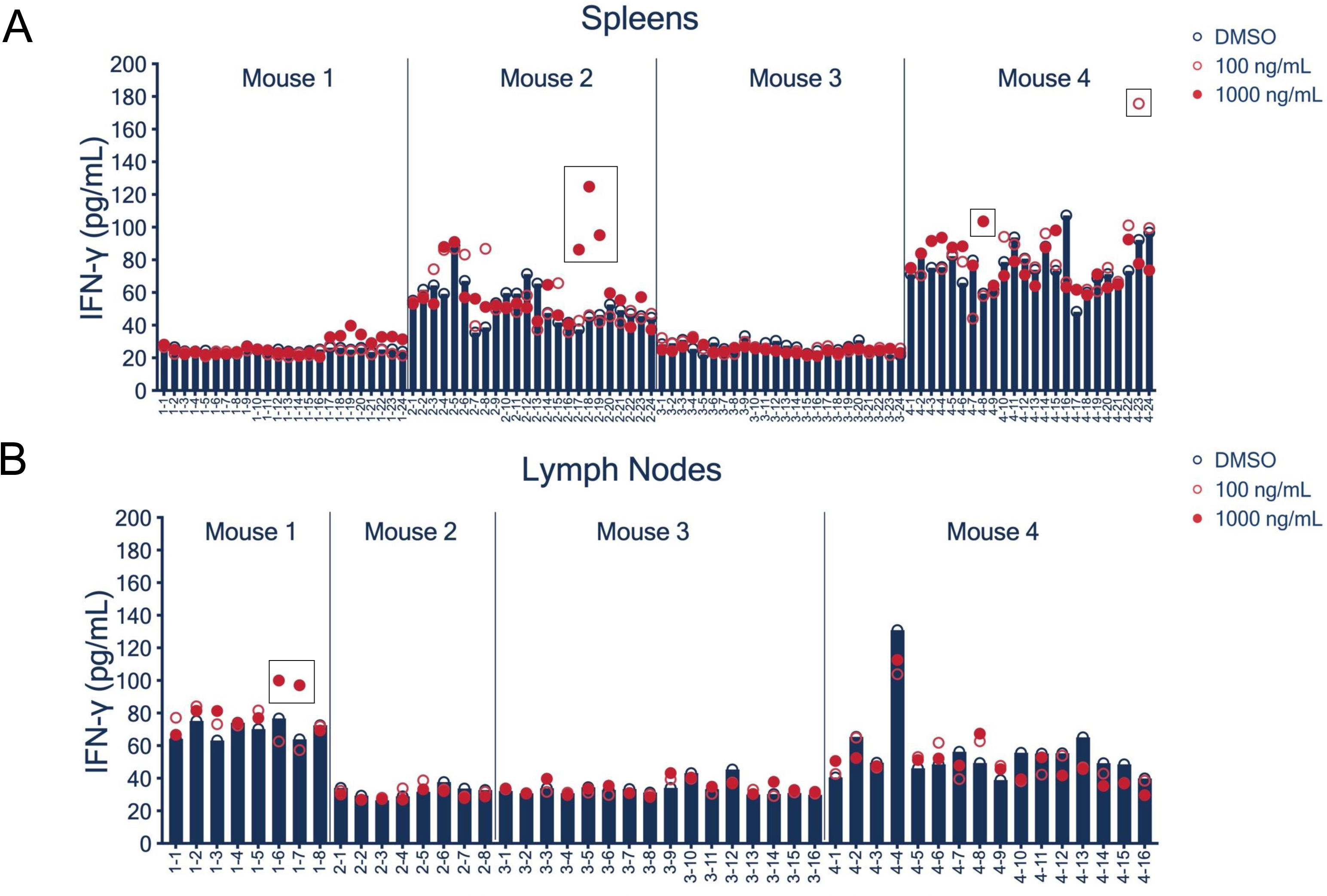
mtNPM1 peptide-reactive T cells were detected in three HLA-A*11:01-transgenic mice vaccinated with AVEEVSLRK. Following the vaccination series, spleens and inguinal lymph nodes from four vaccinated mice were processed into single-cell suspensions and stimulated *in vitro* with AVEEVSLRK in up to 24 individual cultures per tissue. Cells were then co-cultured with HLA-A*11:01-expressing K562 cells pulsed with AVEEVSLRK at 100 or 1,000 ng/mL, or with DMSO as a control. The following day, IFN-γ concentrations in co-culture supernatants were measured by ELISA. IFN-γ production by spleen- and lymph node-derived cultures is shown in panels (A) and (B), respectively. Cultures selected for TCR isolation are indicated by boxed data points.

**Figure 3.**
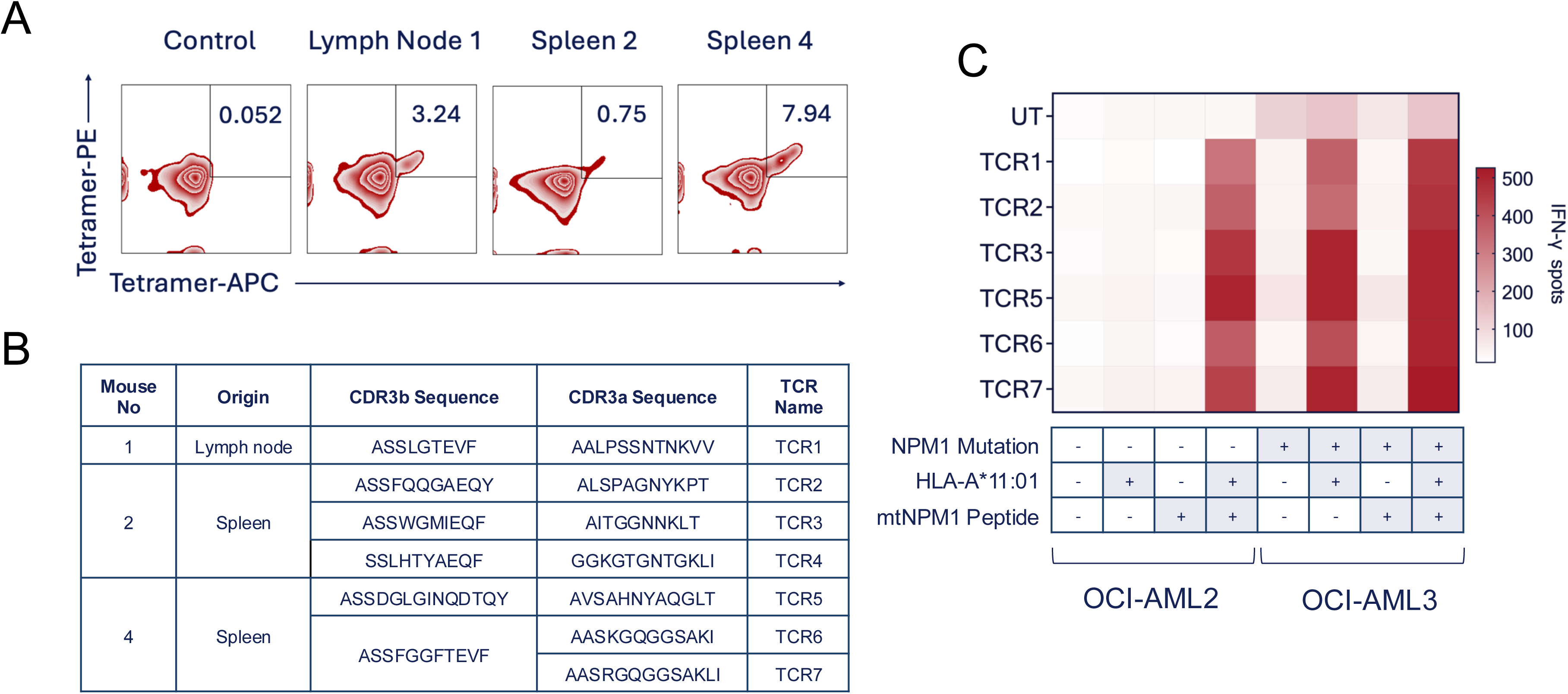
Six functional mtNPM1-reactive TCRs were identified from HLA-A*11:01-transgenic mice vaccinated with AVEEVSLRK. **(A)** Reactive murine lymphocyte cultures shown in Figure 2 were pooled separately for each mouse and stained with HLA-A*11:01-mtNPM1 tetramers. Plots, gated on live CD8^+^ cells, show the percentage of tetramer double-positive cells, which were sorted for single-cell TCR sequencing. Splenocytes from an HLA-A*11:01-negative mouse served as a staining control. **(B)** Seven unique TCRs (TCR1–TCR7) were identified from the populations shown in panel A; the table shows their CDR3α and CDR3β amino acid sequences. TCR6 and TCR7 shared the same β chain but had distinct α chains. **(C)** Human T cells transduced with each TCR were first tested for recognition of the mtNPM1 peptide; six of the seven TCRs showed peptide reactivity (Supplementary Figure 2). T cells bearing these six TCRs were then co-cultured with leukemia cell lines expressing either wild-type NPM1 (OCI-AML2) or mtNPM1 (OCI-AML3), with or without retroviral transduction of HLA-A*11:01, which is not naturally present on either cell line. AML cells were either pulsed with AVEEVSLRK or left unpulsed before co-culture; peptide pulsing served as a control for successful HLA-A*11:01 expression. Heatmap shows the results of IFN-γ ELISpot. The mean of two technical replicates is shown. These findings were confirmed using T cells from another healthy donor (not shown).

Retroviral supernatants were generated using HEK293 cells (ATCC, Manassas, VA) with stable Gag and Pol expression (293GP cells). They were plated in poly-D-lysine-coated 6-well plates (Corning Incorporated) at 1 × 10^6^ cells/well in Dulbecco’s Modified Eagle Medium (DMEM) supplemented with 10% FBS and 1% GlutaMAX. After 6 hours of incubation at 37 °C and 5% CO_2_, cells were co-transfected with 2 µg of MSGV1 plasmid (encoding TCRα, TCRβ, or TCRα/β) and 1 µg of RD114 envelope plasmid using Lipofectamine 2000 (Thermo Fisher Scientific, Waltham, MA). The following day, transfected cells were supplemented with 1 mL of complete DMEM.

To prepare T cells for transduction, healthy donor peripheral blood mononuclear cells (PBMCs), obtained from the National Institutes of Health (NIH) Department of Transfusion Medicine (DTM) by leukapheresis, were cultured for 2 days in complete T-cell medium (RPMI 1640 supplemented with 10% human serum [GeminiBio, West Sacramento, CA], 1% HEPES, GlutaMAX, and Anti-Anti]), together with 50 ng/mL anti-CD3 antibody (OKT3; eBioscience, San Diego, CA) and 300 IU/mL rhIL-2.

For transductions, retroviral supernatants were harvested 2 days after transfection and centrifuged at 300 × *g* for 5 minutes at room temperature to remove cellular debris. These supernatants, consisting either of pooled TCRα and TCRβ supernatants or a single TCRα/β supernatant, were added at 2 mL/well to non-tissue-culture-treated 6-well plates (Corning, Corning, NY) precoated with 0.01 mg/mL RetroNectin (Takara Bio). Plates were centrifuged at 2,000 × *g* at 32 °C for 2 hours, with acceleration and deceleration settings of 4. Following centrifugation, 2 × 10^6^ stimulated PBMCs resuspended in 2 mL of complete T-cell medium supplemented with 30 IU/mL rhIL-2 were added to each well. Plates were then centrifuged at 1,000 × *g* at 25 °C for 10 minutes, with acceleration and deceleration settings of 1, and incubated overnight at 37 °C and 5% CO_2_. The following day, cells were supplemented with 2 mL of complete T-cell medium containing rhIL-2. Transduction efficiency was assessed by flow cytometry based on murine TCRβ expression in human T cells.

For all experiments involving TCR6 and TCR7, except the initial screening experiments, CD8^+^ TCR^+^ cells were purified by cell sorting and expanded (Supplementary Figure 1). For this, transduced CD3^+^ CD8^+^ T cells expressing murine TCRs were first bulk-sorted using the SH800S Cell Sorter. They were then subjected to a rapid expansion protocol by co-culturing them with a >100-fold excess of irradiated (60 Gy) allogeneic PBMCs in 50/50 medium (a 1:1 mixture of complete T-cell medium and AIM-V [Thermo Fisher Scientific] supplemented with 5% human serum and 1% Anti-Anti), together with 30 ng/mL OKT3 and 3,000 IU/mL rhIL-2. A half-medium change was performed on day 7, and culture composition was assessed by flow cytometry on day 12 to 14. Cultures in which CD8^+^ TCR^+^ cells represented >85% of the live-cell population were considered suitable for further testing and were used either fresh or cryopreserved until further use. To determine the CD8 co-receptor dependence of TCR6 and TCR7, the same procedure was performed in parallel for CD4^+^ TCR^+^ cells.

### TCR Cell and Peptide Recognition Assays

HPLC-purified peptides were obtained from GenScript (Piscataway, NJ) as lyophilized powders, resuspended in DMSO at 10 mg/mL, and stored at −20 °C until use. Peptide recognition was evaluated by overnight co-culture of TCR-transduced T cells (2 × 10^4^/well) with peptide-pulsed antigen-presenting cells (1 × 10^5^/well) in 96-well plates, followed by measurement of IFN-γ production as described above. For most experiments, K562-A*11:01 cells served as antigen-presenting cells. They were pulsed with individual peptides at a final concentration of 1 or 10 μg/mL (unless indicated otherwise) at 37 °C overnight, followed by two washes before co-culture. For cell-recognition assays, the same co-culture conditions were used, with K562 cells replaced by the indicated target cells.

### Leukemia Cell Lines

For recognition and cytotoxicity assays, two human AML cell lines were used: OCI-AML2, which expresses wild-type NPM1, and OCI-AML3, which expresses mtNPM1. Both lines were obtained from DSMZ (Leibniz, Germany) and were maintained in Iscove’s Modified Dulbecco’s Medium (IMDM; Thermo Fisher Scientific) supplemented with 20% FBS.

For TCR testing, both cell lines were retrovirally transduced to express HLA-A*11:01 and a puromycin resistance gene using the same protocol described for K562 cells^15^. Three days after transduction, puromycin selection was initiated at 0.001 mg/mL, and was maintained during subsequent culture; puromycin was withdrawn before experiments involving T cells.

For use in IncuCyte cytotoxicity assays (see below), HLA-A*11:01-transduced OCI-AML2 and OCI-AML3 cells were further transduced with a retroviral vector encoding the red fluorescent protein mKate2 fused to a nuclear localization signal. Two weeks after transduction, mKate2-positive populations were purified by cell sorting. mKate2 expression was confirmed by fluorescence microscopy before use in co-culture experiments.

For xenograft experiments, the HLA-A*11:01-expressing OCI-AML3 cell line was transduced with a lentiviral construct encoding firefly luciferase and green fluorescent protein (GFP)^19^. Two weeks after transduction, GFP-positive populations were purified by cell sorting. Luciferase activity was confirmed *in vitro* by adding 10 μl of 15 mg/mL D-luciferin (Revvity, Waltham, MA) solution to 1 × 10^6^ cells in 100 μl of medium, incubating for 5 minutes at room temperature, and measuring luminescence using a SpectraMax iD3 Microplate Reader (Molecular Devices, San Jose, CA).

### Primary AML Patient Samples

Deidentified PBMC samples containing primary AML (pAML) cells with or without HLA-A*11:01 and type A mtNPM1 were obtained from the Stem Cell and Xenograft Core at the Perelman School of Medicine, University of Pennsylvania (RRID: SCR_010035). To minimize cell aggregation after thawing, samples were rapidly thawed in a 37 °C water bath and immediately diluted in 9 mL of 100% FBS before washing. Cells were then cultured in IMDM supplemented with 20% FBS.

A portion of each sample was used to confirm the AML phenotype reported in the clinical record by flow cytometry (see below). Concurrently, DNA was extracted using the DNeasy Blood & Tissue Kit (Qiagen, Hilden, Germany) according to the manufacturer’s instructions. HLA typing was then confirmed by next-generation sequencing at the NIH DTM, and type A NPM1 mutation status was validated by PCR. For the latter, mtNPM1-specific primers were used: forward (spanning the type A mutation site) 5′-GCTATTCAAGATCTCTGTGTGGC-3′; reverse (annealing to the wild type part of the sequence), 5′-TGGACAACACATTCTTGGCA-3′ (Integrated DNA Technologies, Coralville, IA). PCR products were analyzed using the 4200 TapeStation System with D1000 ScreenTape (Agilent Technologies, Santa Clara, CA) according to the manufacturer’s protocol.

### TCR-Mediated Cytotoxicity Assays

TCR-mediated lysis of target cells was analyzed using IncuCyte for leukemia cell lines and flow cytometry for primary AML cells. For the former, 2 ×10^4^ TCR-T cells were co-cultured with 1 ×10^4^ HLA-A*11:01- and mKate2-transduced OCI-AML lines in complete T cell medium in non-tissue-culture-treated 96-well plates (Thermo Fisher Scientific, Waltham, MA) pre-coated with 0.01 mg/mL RetroNectin. For experiments with varied effector-to-target (E:T) ratios, the number of tumor cells was kept constant. Co-cultures were incubated at 37 °C with 5% CO_2_ for 96 hours; images of each well were taken every 4 hours using the Live-Cell Imaging and Analysis System (IncuCyte; Sartorius, Göttingen, Germany). The number of tumor cells expressing mKate2 was quantified using the IncuCyte Cell-by-Cell Analysis Software; counts in each well were normalized to their respective baseline values. Percentage cytotoxicity at each time point was calculated as 100 × [1 − (normalized tumor cell count in the test co-culture / mean normalized tumor cell count in tumor-only controls)]. The same procedure was used in experiments with OCI-AML3-A*11:01 cells transduced with a luciferase-GFP construct, with GFP-positive tumor cells quantified by live-cell imaging.

For primary AML cytotoxicity assays, TCR-T cells were first stained with CellTrace Violet (CTV; Thermo Fisher Scientific) according to the manufacturer’s instructions. CTV-labeled TCR-T cells (2 × 10^4^) were then co-cultured with 1 × 10^4^ pAML cells in 96-well plates (Corning) for 72 hours at 37 °C with 5% CO_2_. Viable tumor cells were quantified by flow cytometry by gating on live, CTV^neg^ CD3^neg^ CD19^neg^ CD14^neg^ CD16^neg^ cells and normalizing their counts to CountBright Absolute Counting Beads (Thermo Fisher Scientific) added immediately before acquisition. For pAML11, which expressed CD14, the gating strategy was adjusted accordingly. Percentage cytotoxicity was calculated as 100 × [1 − (normalized tumor cell count in the test co-culture / mean normalized tumor cell count in tumor-only controls)].

### Peptide Cross-Reactivity Assays

For alanine-substitution assays, a series of peptides was synthesized so that each amino acid in the reference peptide AVEEVSLRK was individually substituted with alanine, except for the native alanine at position 1, which was replaced with glycine. Peptide reactivity was then assessed by pulsing K562-A*11:01 cells and co-culturing them with TCR-T cells, as described above. Residues were considered essential for TCR recognition if their substitution resulted in at least a 90% reduction in IFN-γ concentration compared with the reference peptide. The resulting recognition motifs were queried against the human proteome using the ExPASy ScanProsite tool (https://prosite.expasy.org/scanprosite/).

For cross-reactivity testing, peptides with physicochemical similarity to AVEEVSLRK were identified using amino acid grouping based on physicochemical properties and the BLOSUM62 substitution matrix with a threshold score of ≥1^20^. For the physicochemical grouping strategy, the AVEEVSLRK query sequence was defined as [AGST]-[VILM]-[DENQ]-[DENQ]-[VILM]-[AGST]-[VILM]-[RKH]-[RKH], where brackets indicate the amino acids acceptable at each position. For the BLOSUM62 strategy, the corresponding query sequence was [AS]-[ILMV]-[DEKQ]-[DEKQ]-[ILMV]-[ANST]-[ILMV]-[KQR]-[KEQR]. Both query sequences were searched against the human proteome using the ExPASy ScanProsite tool, generating a list of candidate peptides (Supplementary Figure 6). Binding of these peptides to HLA-A*11:01 was predicted using NetMHCpan-4.1 (https://services.healthtech.dtu.dk/services/NetMHCpan-4.1/); peptides classified as strong or weak predicted binders (defined by ranks of <0.5 and 0.5–2, respectively) were synthesized and tested for recognition by TCR-T cells as described above.

### Assessment of Normal Cell Recognition by TCR-Transduced T Cells

To further assess the cross-reactive potential of the TCRs described in this study, HLA-A*11:01-positive and -negative PBMCs or peripheral blood stem cells (PBSCs) were co-cultured at 1 × 10^5^ cells/well with TCR-T cells (2 × 10^4^ cells/well) in 96-well plates. TCR-T recognition was then assessed as described above. PBMCs were obtained from leukapheresis products from healthy volunteers and isolated by density-gradient centrifugation. Samples were analyzed by flow cytometry to ensure adequate representation of major PBMC populations, including T cells (CD3^+^), B cells (CD19^+^), monocytes (CD14^+^), and NK cells (CD56^+^). HLA typing was performed by next-generation sequencing at the NIH DTM.

To obtain PBSCs, residual leukapheresis samples from HLA-typed granulocyte colony-stimulating factor (G-CSF)- or plerixafor-mobilized HSCT donors were subjected to CD34^+^ cell isolation using the CD34 MicroBead Kit (Miltenyi Biotec, Gaithersburg, MD) according to the manufacturer’s instructions. Sample purity was confirmed by flow cytometry using the anti-CD34 antibody.

To assess the alloreactivity potential of the TCRs, a panel of Epstein-Barr virus-transformed B-cell (EBV-B) lines was generated from HLA-typed healthy donors to provide broad coverage of diverse HLA alleles (see table within Supplementary Figure 6). To do this, up to 1 × 10^8^ PBMCs were cultured in 4 mL of complete RPMI medium supplemented with 1 mL of B95-8 culture supernatant containing EBV (ATCC) and 0.5 μg/mL cyclosporine A (Sigma-Aldrich, St. Louis, MO) for 4–6 weeks. Medium containing cyclosporine A was replenished as needed. Cultures were validated by flow cytometry for CD19 positivity and absence of CD3 expression, after which cells were further expanded in cyclosporine-free medium. They were then co-cultured with TCR-T cells under the same conditions described above.

### AML Xenograft Studies

Female 4-to 6-week-old NOD.Cg-Prkdc^scid^ Il2rg^tm1Wjl^/SzJ (NSG) mice (The Jackson Laboratory, Bar Harbor, ME) were administered 1 × 10^6^ OCI-AML3-A*11:01 cells expressing a GFP-luciferase construct by tail-vein injection on day 0. On day 6 (experiment shown in Figure 7) or day 3 (experiment shown in Supplementary Figure 7), 10 × 10^6^ CD8^+^ TCR^+^ T cells expressing TCR6, TCR7, or an irrelevant TCR were administered via the same route; PBS injections served as an additional negative control. Tumor engraftment and growth were assessed by serial bioluminescence imaging after intraperitoneal injection of 3 mg D-luciferin (Caliper Life Sciences, Hopkinton, MA) using the IVIS Lumina Imaging System (PerkinElmer). Luminescence was quantified using the RadianceQuantifier graphical user interface. On day 35, mice were euthanized and their bone marrow, liver, spleen, and extramedullary tumors were harvested for flow cytometric analysis of tumor and T cells.

**Figure 4.**
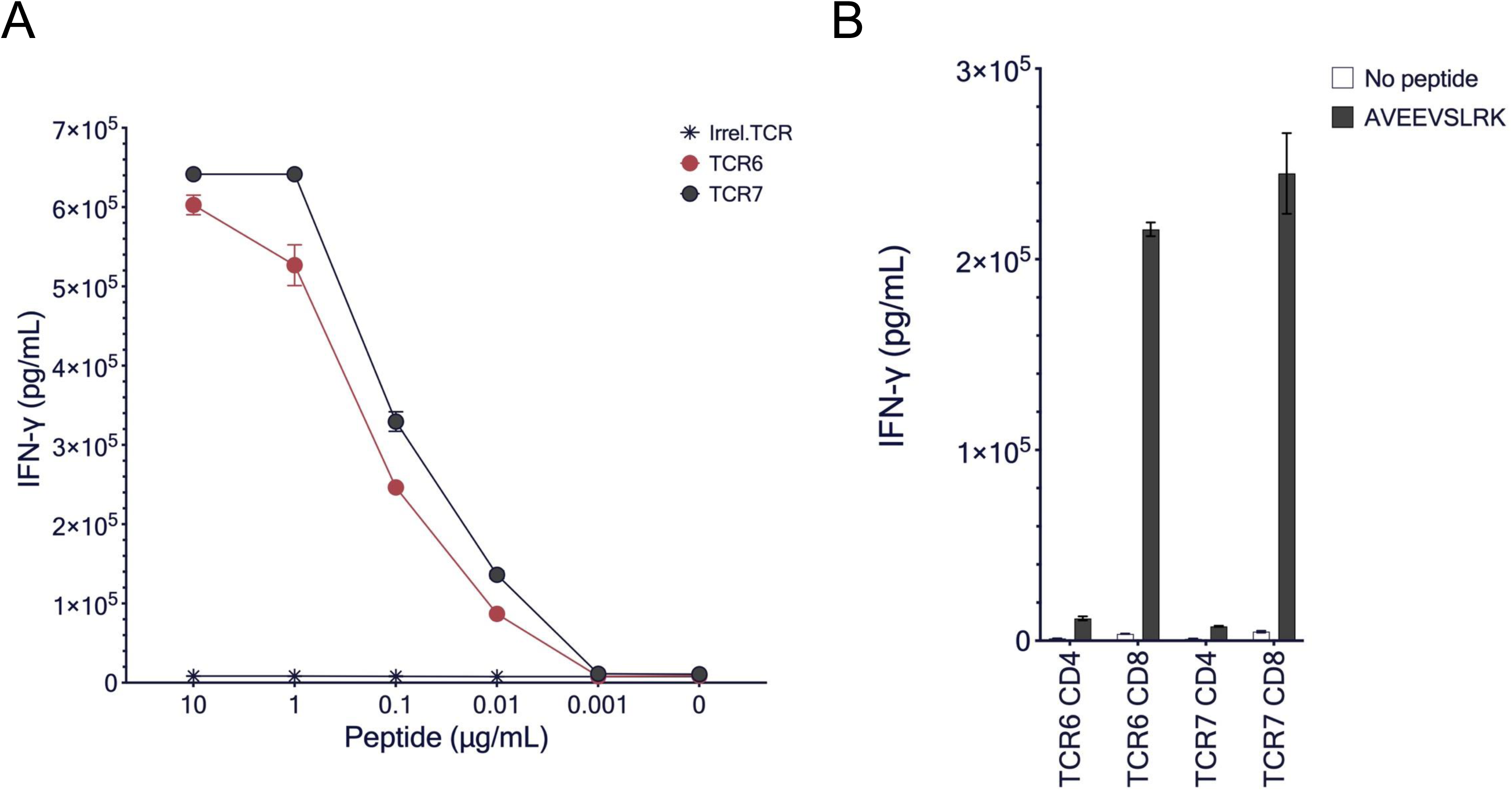
TCR6 and TCR7 recognize AVEEVSLRK in a dose-dependent and co-receptor-dependent manner. **(A)** Human T cells were transduced with TCR6, TCR7, or an irrelevant TCR and co-cultured overnight with HLA-A*11:01-overexpressing K562 cells pulsed with decreasing concentrations of AVEEVSLRK. Line graph shows IFN-γ production measured using the IFN-γ U-PLEX assay. **(B)** TCR6- and TCR7-transduced human T cells were sorted into CD4^+^ TCR^+^ and CD8^+^ TCR^+^ populations and co-cultured overnight with K562-A*11:01 cells pulsed with AVEEVSLRK (1 µg/mL). Bar graph shows IFN-γ production measured using the U-PLEX assay. For both panels, the mean and standard deviation of two technical replicates are shown for each condition. These findings were confirmed using T cells from unrelated healthy donors (Supplementary Figure 3).

**Figure 5.**
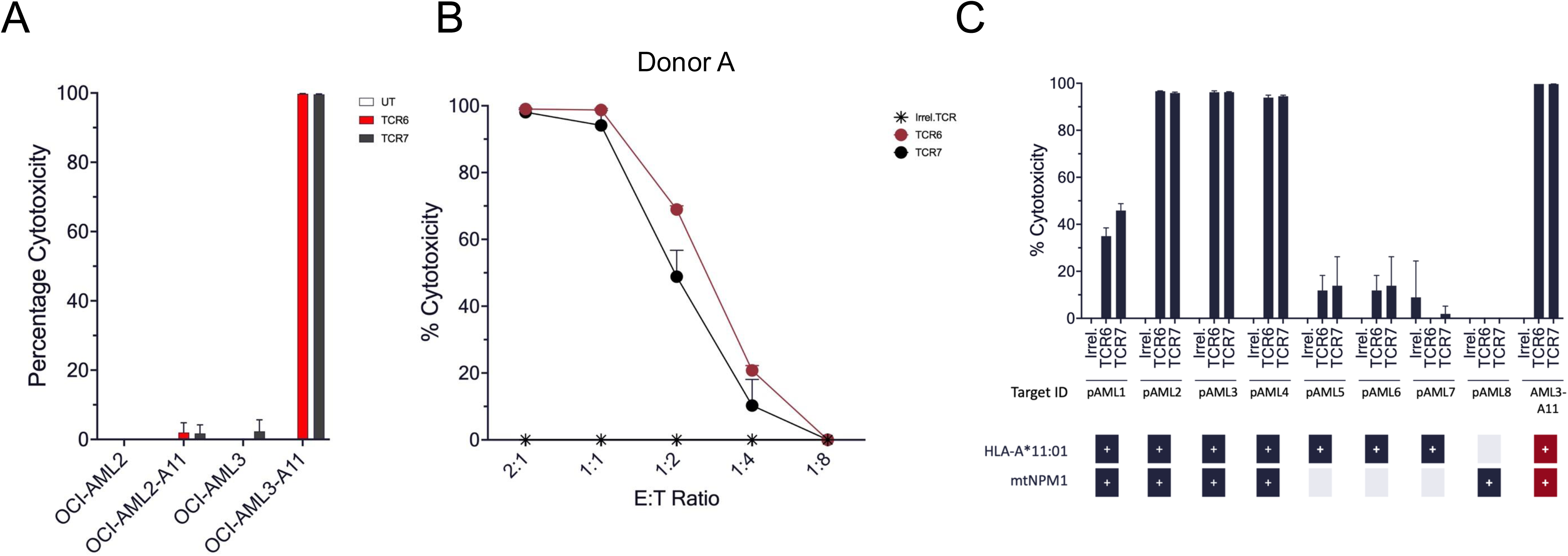
TCR6 and TCR7 mediate specific and robust *in vitro* lysis of leukemia cells co-expressing HLA-A*11:01 and mtNPM1. **(A)** OCI-AML2 (wild-type NPM1) and OCI-AML3 (mtNPM1) cells, with or without HLA-A*11:01 expression, were engineered to express mKate2, a red fluorescent marker. They were then co-cultured with TCR6- or TCR7-transduced T cells for 96 hours, with tumor cells imaged in real time and quantified using the IncuCyte Live-Cell Analysis System. Negative controls included tumor-only wells and co-cultures with untransduced (UT) T cells. Bar graph shows percentage cytotoxicity after 96 hours of co-culture, calculated as (1 – normalized test co-culture cell count / mean normalized tumor-only cell count) × 100. The full time course and data using T cells from another healthy donor are shown in Supplementary Figure 4. **(B)** The experiment in panel A was repeated using OCI-AML3-HLA-A*11:01 cells co-cultured with T cells transduced with TCR6, TCR7, or an irrelevant TCR at decreasing effector-to-target (E:T) ratios. Line graph shows percentage cytotoxicity at 96 hours. A representative experiment showing the lower range of cytotoxic activity is shown; representatives of experiments showing greater activity are presented in Supplementary Figure 4B. **(C)** TCR-T cells were co-cultured at a 2:1 E:T ratio with a panel of 8 primary AML samples (pAML1–8) or OCI-AML3-HLA-A*11:01 cells (labeled AML3-A11) as a positive control. Bar graph shows percentage cytotoxicity at 72 hours, calculated based on the number of viable AML cells obtained by flow cytometry using the same formula as in panel A. The table below the graph shows the HLA-A*11:01 and mtNPM1 status of each sample. For panels (A) and (B), mean ± SD of two technical replicates is shown; for panel (C), three technical replicates. Results were confirmed in two independent experiments shown in Supplementary Figure 5.

**Figure 6.**
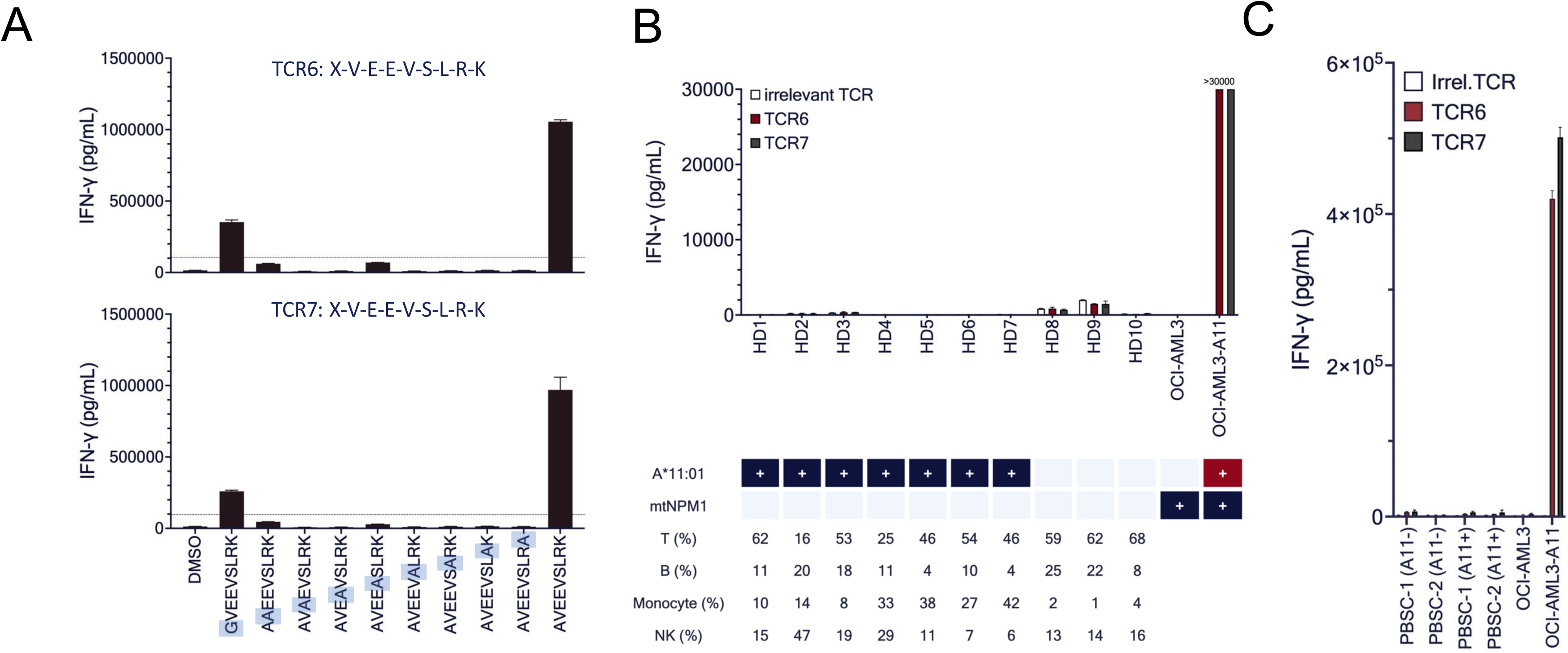
TCR6 and TCR7 demonstrated favorable *in vitro* safety profiles in peptide- and cell-based assays. **(A)** TCR6- and TCR7-transduced T cells were co-cultured with K562-A*11:01 cells pulsed with DMSO (negative control), 1 µM AVEEVSLRK (reference peptide), or equal concentrations of peptides in which individual amino acids were serially substituted with alanine (or glycine, when the original residue was alanine). Bar graphs show IFN-γ secretion by TCR6 (top) and TCR7 (bottom), measured the following day using the IFN-γ U-PLEX assay. Residues were considered essential for TCR recognition if their substitution resulted in at least a 90% reduction in IFN-γ secretion compared with AVEEVSLRK. This analysis defined the recognition motif for both TCR6 and TCR7 as X-V-E-E-V-S-L-R-K, where X represents any amino acid. When queried against the human proteome using the ExPASy ScanProsite tool, this motif yielded no matches. Results were confirmed using T cells from an independent healthy donor (not shown). **(B)** T cells transduced with TCR6, TCR7, or an irrelevant TCR were co-cultured overnight with peripheral blood mononuclear cells (PBMCs) from multiple HLA-A*11:01-positive or -negative healthy donors. Bar graph shows IFN-γ production measured by U-PLEX. The table below shows the T-cell, B-cell, monocyte, and NK-cell composition of each PBMC sample, as determined by flow cytometry. OCI-AML3 cells lacking HLA-A*11:01 (AML3) and OCI-AML3 cells transduced with HLA-A*11:01 (AML3-A11) served as negative and positive controls, respectively. Results were confirmed using T cells from an unrelated donor (not shown). **(C)** CD34^+^ cells were isolated from peripheral blood stem cell products obtained from two HLA-A*11:01-negative and two HLA-A*11:01-positive stem cell transplant donors and co-cultured overnight with T cells transduced with TCR6, TCR7, or an irrelevant TCR. Bar graph shows IFN-γ production measured by U-PLEX. The same negative and positive controls described in panel B were used. For all panels, mean ± SD of 2–4 technical replicates is shown.

**Figure 7.**
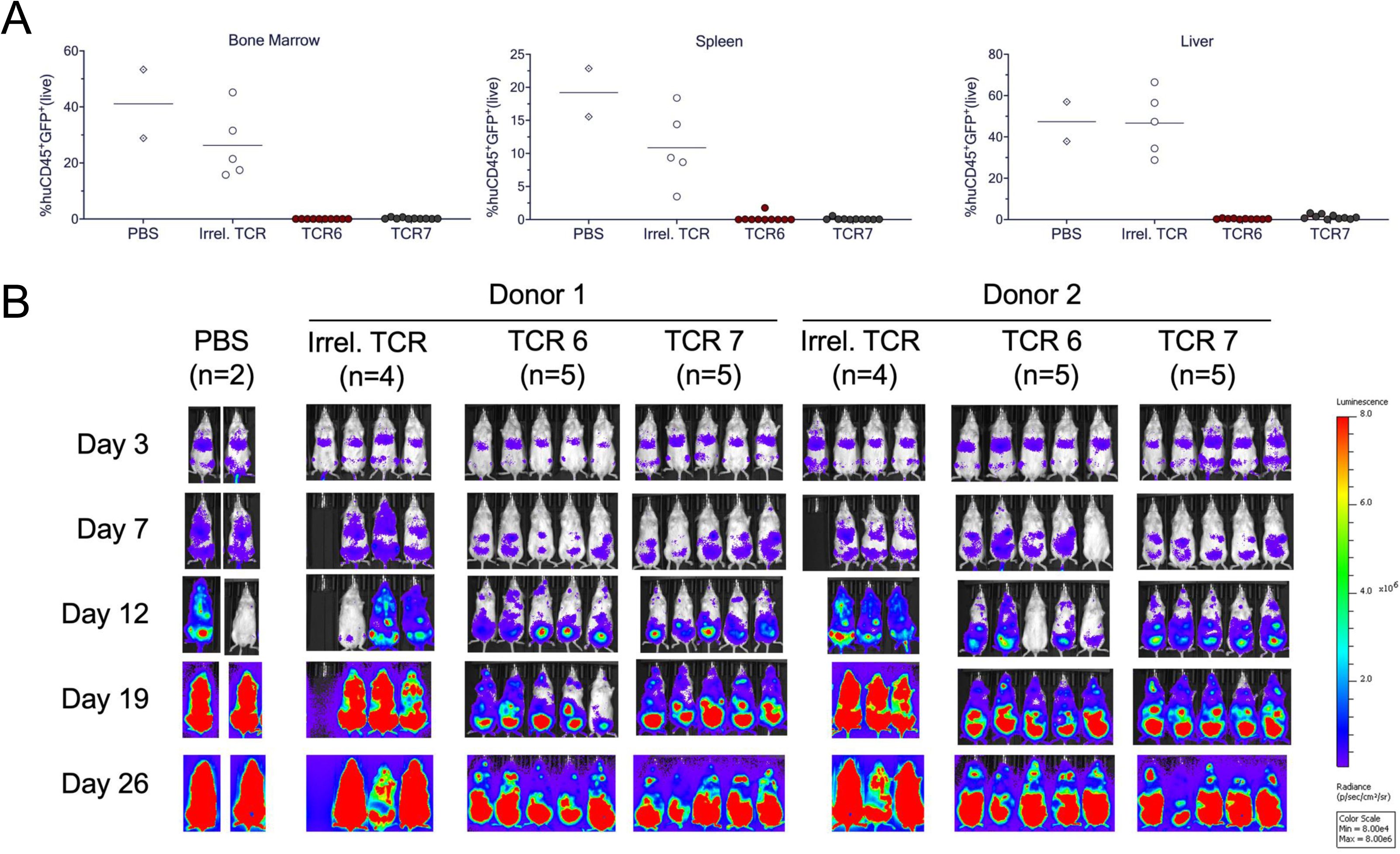
Adoptive transfer of TCR6- and TCR7-transduced T cells decreased AML burden and achieved near-complete clearance of leukemia from the hematopoietic compartments of NSG mice. (A) On day 0, 1 × 10^6^ OCI-AML3 cells engineered to express HLA-A*11:01 and a GFP-luciferase construct were injected into 4-week-old female NSG mice via tail veins. On day 6, 10 × 10^6^ CD8^+^ TCR^+^ T cells from two healthy donors, transduced with TCR6, TCR7, or an irrelevant TCR, were administered by tail vein injection. On day 35, all mice were sacrificed, and spleens, livers, and bone marrow samples were collected and analyzed by flow cytometry. Plots show the percentage of human CD45^+^ GFP^+^ AML cells among live cells for each treatment group. Each symbol represents an individual mouse, and lines indicate mean values. For this analysis, data from the two donors were pooled; one mouse from the donor 2 irrelevant TCR group could not be analyzed. (B) Starting before T-cell infusion, leukemia progression was monitored by serial bioluminescence imaging.

### Statistical Analysis

Data analysis and graphing were performed using GraphPad Prism 10.2 (GraphPad Software, La Jolla, CA). As this study was primarily descriptive, no formal statistical hypothesis testing was performed. When applicable, data are presented as mean ± standard deviation (SD).

## Results

### Six Functional mtNPM1- and HLA-A*11:01-Specific TCRs Were Identified from HLA-A*11:01-Transgenic Mice Vaccinated with AVEEVSLRK

Four HLA-A*11:01-transgenic mice were vaccinated with the mtNPM1-derived 9-mer peptide AVEEVSLRK. One week after the final vaccination, lymphocytes isolated from their spleens and lymph nodes were stimulated *in vitro* with AVEEVSLRK to enrich potentially reactive T cells. Individual T-cell cultures were then tested for recognition of peptide-pulsed K562-A*11:01 cells. As shown in Figure 2, peptide-reactive T cells were detected in three mice (lymph node-derived cells from mouse #1 and spleen-derived cells from mouse #2 and #4), based on increased IFN-γ production relative to the negative control.

Reactive cells derived from the same mouse were pooled and stained with HLA-A*11:01-mtNPM1 tetramers. Tetramer-positive cells were detected in and sorted from all three pooled cultures (Figure 3A). Single-cell TCR sequencing of the sorted populations identified seven unique TCRs (Figure 3B). Their α and β chains were cloned into a retroviral expression vector and subsequently transduced into human T cells for downstream testing (Supplementary Figure 1).

Following expression in human T cells, six of the seven murine TCRs recognized AVEEVSLRK-pulsed K562-A*11:01 cells (Supplementary Figure 2). All six also induced IFN-γ production in response to the OCI-AML3 cells, which naturally express mtNPM1 and were transduced to express HLA-A*11:01. In contrast, they did not recognize OCI-AML3 cells lacking HLA-A*11:01 or the wild-type NPM1 OCI-AML2 cell line, regardless of whether it was transduced with HLA-A*11:01 (Figure 3C).

These results demonstrated that the six TCRs recognized endogenously processed mtNPM1 in an HLA-A*11:01-restricted manner. In subsequent experiments, all six functional TCRs were evaluated in parallel, and two receptors (TCR6 and TCR7) consistently mediated specific lysis of AML cell lines and primary AML samples. All further characterization therefore focused on these receptors, which shared the same TCR β chain and had closely related α chains.

### TCR6 and TCR7 Recognized mtNPM1 in a Dose-Dependent and CD8 Co-receptor-Dependent Manner

To assess functional avidity, human CD8^+^ T cells from two healthy donors were transduced with TCR6, TCR7, or an irrelevant control TCR and co-cultured with K562-A*11:01 cells pulsed with decreasing concentrations of AVEEVSLRK. Only T cells expressing TCR6 or TCR7 produced comparable, dose-dependent levels of IFN-γ, with reactivity above background at mtNPM1 peptide concentrations as low as 10 ng/mL (Figure 4A and Supplementary Figure 3A).

Because TCR6 and TCR7 differed by three amino acids in their α chains, including one residue within the CDR3α region, three mutant receptors were generated in which each of these residues was individually substituted with alanine to assess their functional relevance. Recognition of K562-A*11:01 cells pulsed with AVEEVSLRK by these mutant receptors was then compared with that of the unmodified TCR6 and TCR7. Only the substitution of the CDR3α residue abolished activity (Supplementary Figure 3B), suggesting that this differing residue is functionally important and supporting TCR6 and TCR7 as closely related receptors.

To determine whether recognition required a T-cell co-receptor, transduced cells were simultaneously FACS-sorted into TCR^+^ CD8^+^ and TCR^+^ CD4^+^ populations. While both TCR6- and TCR7-transduced CD8^+^ T cells produced IFN-γ in response to AVEEVSLRK-pulsed K562-A*11:01 cells, the corresponding CD4^+^ populations showed minimal response (Figure 4B and Supplementary Figure 3C). Together, these findings demonstrate that both receptors recognize mtNPM1 in a peptide dose-dependent manner and require the CD8 co-receptor for activity.

### TCR6 and TCR7 Mediated Robust and Specific Lysis of mtNPM1-Positive, HLA-A*11:01-Positive Leukemia Cells

Next, TCR6 and TCR7 were tested for their ability to specifically lyse the HLA-A*11:01-expressing OCI-AML3 cell line. CD8^+^ TCR-T cells were co-cultured with fluorescently labeled OCI-AML3 cells or, as a wild-type NPM1 control, OCI-AML2 cells, with or without prior HLA-A*11:01 transduction for both lines. Target-cell lysis was measured over 96 hours using a live-cell imaging cytotoxicity assay. As shown in Figure 5A and Supplementary Figure 4, TCR6 and TCR7 mediated complete lysis only of HLA-A*11:01-transduced OCI-AML3 cells, again demonstrating specificity for both mtNPM1 and HLA-A*11:01.

To assess their functional potency, CD8^+^ T cells expressing TCR6, TCR7, or an irrelevant TCR were co-cultured with fluorescently labeled OCI-AML3–HLA-A*11:01 cells at decreasing effector-to-target (E:T) ratios. Both TCR6 and TCR7 maintained cytotoxic activity at E:T ratios ranging from 1:4 to 1:16 across experiments (Figure 5B and Supplementary Figure 4B). Differences in potency between experiments may reflect biological variability between donor T cells or between TCR-T cell preparations.

To test whether the TCRs could lyse primary AML cells with endogenous mtNPM1 and HLA-A*11:01, we obtained corresponding patient samples and controls lacking either mtNPM1 or HLA-A*11:01. In flow cytometry-based cytotoxicity assays, both TCR6 and TCR7 specifically lysed these samples without increased lysis of controls (Figure 5C and Supplementary Figure 5). Because mtNPM1 and HLA-A*11:01 expression could not be reliably quantified, it remains unclear whether lower expression of either antigen or HLA contributed to the reduced cytotoxicity observed against pAML1. Overall, these findings demonstrate that both TCRs can lyse not only an mtNPM1-positive cell line engineered to express HLA-A*11:01, but also unmanipulated primary AML cells obtained from patients with mtNPM1- and HLA-A*11:01-positive disease.

### TCR6 and TCR7 Exhibited Favorable *In Vitro* Safety Profiles

Because these TCRs are of murine origin and did not undergo natural human thymic selection, unexpected cross-reactivity with other human peptides represents an important concern. To assess potential peptide cross-reactivity, motif-based and motif-independent searches were performed. For the motif-based search, TCR recognition motifs were first determined using an alanine substitution assay. A peptide library was generated in which each amino acid in AVEEVSLRK was individually replaced with alanine, or with glycine when the original residue was alanine. These peptides were then pulsed onto K562-A*11:01 cells and tested for recognition by both TCRs. Based on IFN-γ production, TCR6 and TCR7 shared the recognition motif X-V-E-E-V-S-L-R-K, where X represents any amino acid (Figure 6A). A search of the human proteome identified no additional peptides matching this motif.

For the motif-independent search, 14 human peptides with physicochemical properties similar to those of AVEEVSLRK and predicted to bind HLA-A*11:01 were identified as described in the Methods and Supplementary Figure 6A. Neither TCR demonstrated detectable recognition of the tested peptides based on IFN-γ production (Supplementary Figure 6A). Together, these searches suggested a low risk of overt cross-reactivity with other human peptides.

To evaluate recognition of normal, non-tumor cells with physiologic HLA-A*11:01 expression, both TCRs were first tested against a panel of PBMC samples from HLA-A*11:01-positive healthy donors, which contained variable proportions of T cells, B cells, monocytes, and natural killer (NK) cells. Neither TCR induced substantial IFN-γ production in response to any of the samples tested (Figure 6B). The TCRs were next tested for recognition of peripheral blood stem/progenitor cells (PBSCs) obtained by leukapheresis from HLA-A*11:01-positive and - negative HSCT donors, followed by CD34^+^ cell isolation using magnetic beads. As shown in Figure 6C, neither TCR recognized purified HLA-A*11:01-positive PBSCs. Together, these results revealed no detectable cross-reactivity against the normal cells included in our testing panels.

Finally, potential alloreactivity was evaluated using a panel of EBV-immortalized B-cell lines from healthy donors expressing diverse common class I HLA alleles. Neither TCR demonstrated substantial recognition of any of the cell lines tested (Supplementary Figure 6B), providing no evidence of alloreactivity within the evaluated panel.

### TCR6 and TCR7 Controlled Leukemia in the Hematopoietic Compartments *In Vivo*

To evaluate the antileukemic activity of TCR6 and TCR7 *in vivo*, NSG mice were intravenously injected with OCI-AML3 cells engineered to express HLA-A*11:01, GFP, and luciferase. On day 6, mice received CD8^+^ T cells from one of the two healthy donors, engineered to express TCR6, TCR7, or an irrelevant TCR. Leukemia progression was monitored by serial bioluminescence imaging, and tissue leukemia burden was assessed by flow cytometry at sacrifice on day 35.

At sacrifice, flow cytometric analysis demonstrated near-complete clearance of human CD45^+^ GFP^+^ leukemia cells from the bone marrow, spleen, and liver of mice treated with either TCR6 or TCR7 (Figure 7A). A separate experiment in which T cells were administered on day 3 yielded similar results (Supplementary Figure 7A). Nevertheless, all mice developed extramedullary tumors in the lower body. Consistent with this pattern, the distribution of bioluminescence in TCR6- and TCR7-treated mice shifted from systemic disease to localized signal in the lower body, whereas mice receiving PBS or irrelevant TCR-T cells developed progressive disseminated leukemia (Figure 7B and Supplementary Figure 7B). AML cells isolated from these tumors could be eliminated by TCR6 in subsequent *in vitro* co-cultures, indicating retention of sufficient mtNPM1 and HLA-A*11:01 expression for TCR-mediated recognition and killing (Supplementary Figure 8). Thus, extramedullary tumor growth did not appear to be driven by loss of the target antigen or restricting HLA allele. No human T cells were detected in the tumors (data not shown), suggesting that the tumor growth may instead reflect impaired T-cell trafficking or persistence within these lesions. Furthermore, the development of these tumors may reflect the aggressive and atypical growth pattern of the engineered OCI-AML3 model^21,22^, and it thus remains uncertain whether or to what extent this finding is clinically relevant. In summary, these results demonstrate that both TCRs mediated substantial antileukemic activity *in vivo*, including marked control of leukemia in major hematopoietic and visceral compartments.

## Discussion

In this study, we identified two mtNPM1-reactive, HLA-A*11:01-restricted TCRs from HLA-A*11:01-transgenic mice vaccinated with the mtNPM1-derived peptide AVEEVSLRK. These closely related receptors demonstrated comparable activity, specifically recognizing and lysing AML cells expressing both mtNPM1 and HLA-A*11:01. Neither TCR recognized similar human peptides, normal HLA-matched cells, or HLA-mismatched target cells evaluated in this study. In a xenograft model, both receptors mediated near-complete leukemia clearance from the bone marrow, spleen, and liver. Nevertheless, extramedullary tumors developed, potentially reflecting the aggressive growth pattern of the cell-line model used, as discussed below.

For many patients with relapsed or refractory AML, allo-HSCT represents the only potentially curative treatment. However, its use is limited by patient fitness, treatment-related toxicity, and donor availability, leaving many patients ineligible. Its clinical efficacy depends in part on the graft-versus-leukemia (GVL) effect, in which donor T cells recognize antigens such as minor histocompatibility antigens (MiHAs) expressed by recipient AML cells^23,24^. The GVL effect provides a strong rationale for T-cell therapy in AML, while the toxicity resulting from donor T cell recognition of normal tissues, manifested as graft-versus-host disease (GVHD), supports the development of approaches directing T cells toward defined leukemia antigens.

This approach has already been evaluated in AML using engineered T cells. In high-risk patients who had no measurable disease following allo-HSCT, TCR-T targeting WT1 demonstrated encouraging activity in preventing relapse, although it did not induce objective responses in patients with active AML^8,25^. Furthermore, a small study of TCR-T targeting the MiHA HA-1 reported preliminary evidence of antileukemic activity in patients with post-HSCT relapse^26^. Together, these studies establish the feasibility of TCR-T in AML while suggesting that antigen selection and disease burden may influence efficacy.

Although successful neoantigen-directed TCR-T has not yet been reported in AML, recurrent neoantigens derived from driver mutations may offer several advantages over TAAs such as WT1^9^. They are absent from normal tissues, reducing the risk of on-target toxicity. They are also shared, often clonal, and functionally important for maintenance of the malignant phenotype, potentially limiting immune escape through antigen loss.

Mutations in NPM1 exhibit several of these features. They occur in up to one-third of newly diagnosed AML cases, are typically clonal, and are retained at relapse in most patients^27–29^. Although NPM1-mutated AML generally has a favorable prognosis at diagnosis, relapse occurs in up to 50% of patients^30^. Accordingly, NPM1 mutations accounted for 12% of relapsed or refractory AML cases in a large retrospective cohort, with NPM1 mutational status losing its prognostic advantage^27^. Furthermore, although recent trials have shown high response rates with azacitidine, venetoclax, and a menin inhibitor in NPM1-mutated AML, long-term response durability remains uncertain, supporting exploration of TCR-T as a complementary strategy to deepen and sustain remission^31,32^.

The shared frameshifted sequence generated by common NPM1 mutation types produces peptides that are naturally presented by several class I HLA molecules on primary AML cells^13,33^, a critical requirement for TCR-mediated targeting. Spontaneous CD4^+^ and CD8^+^ T-cell responses against these peptides have been detected in patients with NPM1-mutated AML, although responses identified using the same approach in healthy donors make their physiological relevance uncertain^34^. Furthermore, the underrepresentation of several HLA class I alleles capable of presenting mtNPM1-derived peptides among patients with NPM1-mutated AML provides indirect evidence of potential endogenous immune surveillance^35^. However, to our knowledge, no functionally validated mtNPM1-reactive TCR has been isolated from patients, highlighting the need to identify such receptors from alternative sources, including HLA-matched healthy donors or, as demonstrated here, HLA-transgenic mice.

HLA-A*11:01 is carried by approximately 11.2% of the overall US population and is more prevalent in Asian populations, with an estimated carrier frequency of approximately 38% in a large Chinese donor registry^36,37^. Given that NPM1 mutations account for 12% of relapsed or refractory AML cases, and assuming a similar HLA-A*11:01 distribution among patients with AML and the general population, the receptors described in this study could theoretically be applicable to an estimated 1.3% of all patients with relapsed or refractory AML in the United States and 4.5% in China. Although loss or marked downregulation of class I HLA expression is uncommon in AML overall^38,39^, it can occur under post-transplant immune pressure^40,41^, so HLA-A*11:01 expression would need to be confirmed before treatment.

A previously reported HLA-A*11:01-restricted TCR, identified from an HLA-A*11:01-positive healthy donor, recognized the same AVEEVSLRK epitope and mediated specific lysis of primary AML cells *in vitro*^12^. However, this TCR failed to control leukemia in an immunodeficient mouse model designed to avoid the development of extramedullary tumors. Although the apparently greater antileukemic activity observed with our TCRs is encouraging, a head-to-head comparison in the same model would be required to establish superior efficacy.

The murine origin of our TCRs raises two concerns for clinical use. First, murine TCRs have not undergone thymic selection against the human proteome and could therefore exhibit unintended off-target reactivity. However, such reactivity was not identified in our peptide- and cell-based studies, supporting the preliminary safety of both TCRs. Furthermore, reported toxicities in clinical trials using other murine TCRs have resulted from recognition of target antigens or homologous proteins expressed in normal tissues, rather than unexpected recognition of unrelated proteins^42–44^. Second, the potential immunogenicity of murine TCR sequences could limit persistence or repeated administration of TCR-T cells. Although anti-murine immune responses have been reported, they have not been consistently associated with reduced activity^16^. Nevertheless, such responses should be monitored during future clinical testing.

An important limitation of this study is the use of OCI-AML3 cells engineered to express HLA-A*11:01 because no commercially available AML cell line naturally co-expresses mtNPM1 and this HLA. Consequently, HLA-A*11:01 expression in these cells may not fully recapitulate that on patient AML cells. Additionally, OCI-AML3 can exhibit aggressive extramedullary growth in immunodeficient xenograft models, potentially attributable to cell line-specific features such as DNMT3A^R882C^ mutation^21,22^. Although extramedullary disease occurs in patients with NPM1-mutated AML, the extent to which this xenograft model recapitulates patient disease remains uncertain.

In our experiments, TCR6 and TCR7 did not clear the large extramedullary pelvic tumors, whose precise anatomical origin could not be determined because extensive growth obscured the underlying anatomy. Tumor cells isolated from these lesions retained sufficient target antigen and HLA expression for TCR-mediated killing, and no human T cells were detected within the tumors. However, T cells were also undetectable in the bone marrow and spleen, where leukemia had been controlled, limiting interpretation of this finding. Nevertheless, clearance of AML from hematopoietic compartments supports the antileukemic activity of these TCRs in vivo.

To overcome these limitations, OCI-AML3 cells could be selected through serial passage in murine bone marrow to enrich for clones that do not form extramedullary tumors, as previously reported^12^. Alternatively, a patient-derived xenograft model would more closely recapitulate primary AML biology, including more physiologic levels of HLA expression on the tumor cells.

Other study limitations include testing a limited number of primary AML samples, without systematically examining how mtNPM1 and HLA-A*11:01 expression levels influence TCR-mediated cytotoxicity. In addition, the safety analysis was restricted to the peptides and normal cell types tested and therefore cannot exclude recognition of all potential human peptides or tissues. Additional testing using broader panels of primary AML samples and HLA-A*11:01-positive primary cells representing diverse tissues would further strengthen our findings.

In conclusion, we identified two closely related mtNPM1-reactive, HLA-A*11:01-restricted TCRs that demonstrated strong antileukemic activity *in vitro* and within hematopoietic compartments *in vivo*, along with favorable preclinical safety profiles. These findings support their further development for patients with mtNPM1-positive, HLA-A*11:01-positive AML, particularly those with relapsed or refractory disease, for whom effective treatment options remain limited. They also support the use of HLA-transgenic mice as an alternative source of potentially therapeutic TCRs when suitable receptors are difficult to isolate from healthy donors or patients.

## Acknowledgements

This work was supported in part by the Intramural Research Program of the National Institutes of Health (NIH). The contributions of the NIH authors were made as part of their official duties as NIH federal employees, are in compliance with agency policy requirements, and are considered Works of the United States Government. However, the findings and conclusions presented in this paper are those of the authors and do not necessarily reflect the views of the NIH or the U.S. Department of Health and Human Services.

---

**Supplementary Figure 1.**
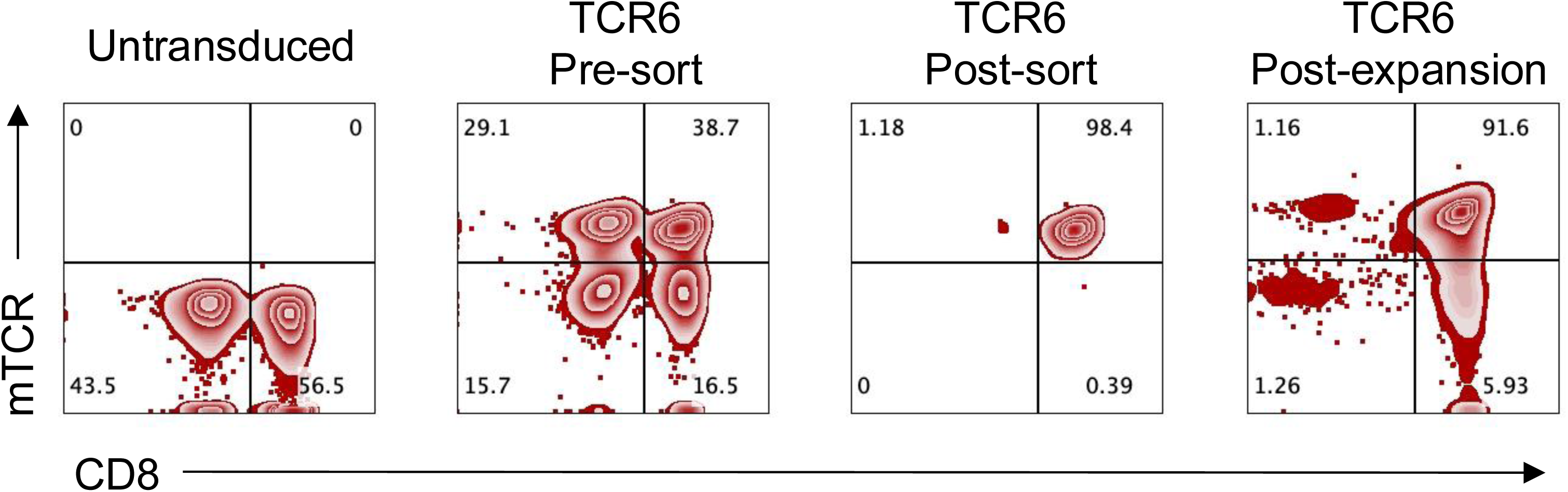
Representative workflow for TCR-T generation. Murine TCRs were retrovirally transduced into peripheral blood T cells from healthy donors for functional testing, followed by sorting for CD8^+^ and mouse TCR^+^ (mTCR^+^) cells and expansion using a rapid expansion protocol (REP). Representative plots show transduction efficiency before and after sorting and REP for TCR6 in one healthy donor; plots are gated on live cells. In most experiments, the final T-cell cultures used for functional testing consisted of >90% CD8^+^mTCR^+^ cells.

**Supplementary Figure 2.**
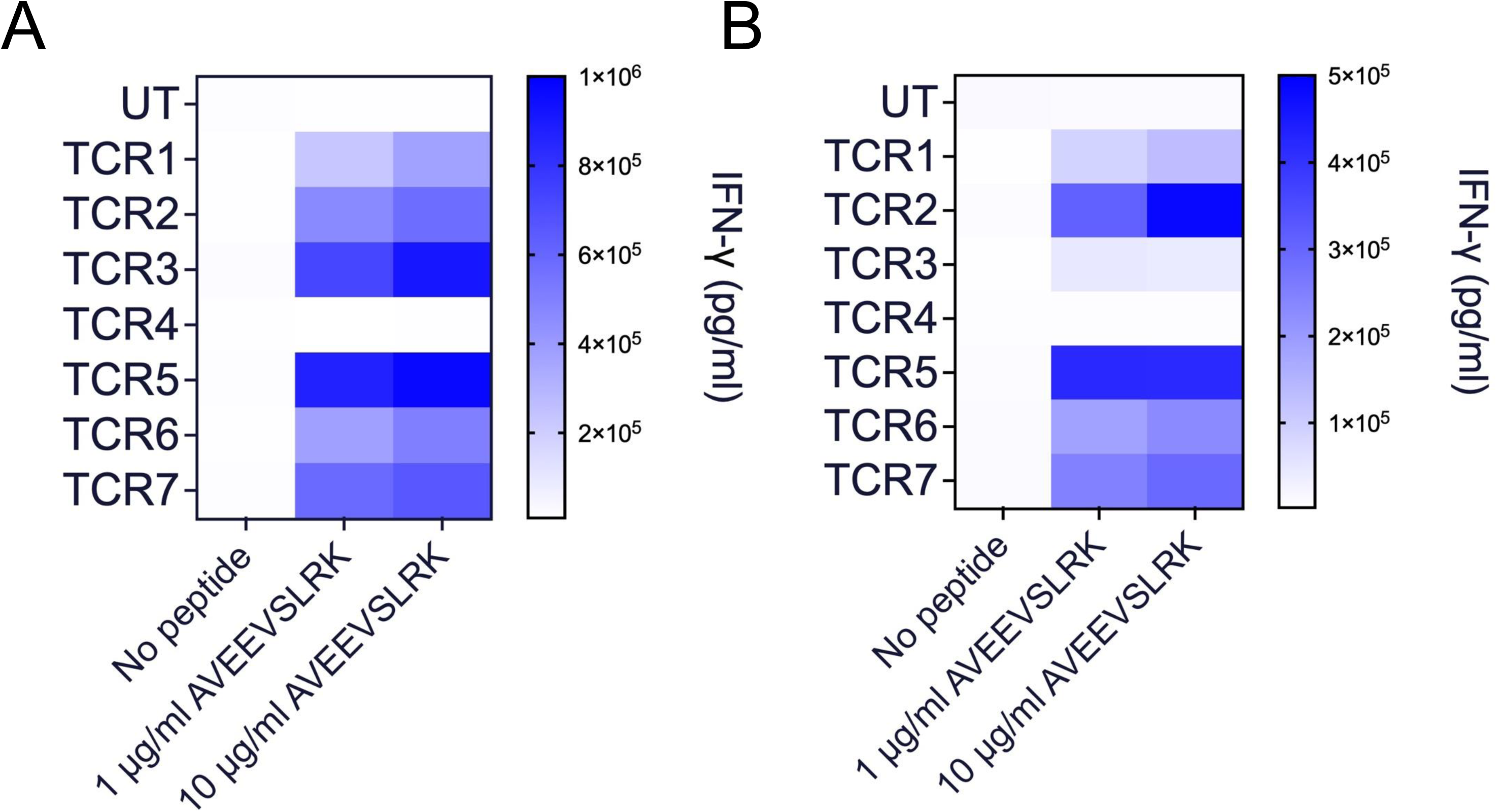
Initial screening of murine TCRs for recognition of the mtNPM1 peptide AVEEVSLRK. Human T cells transduced with TCR1–TCR7 (described in Figure 3) were co-cultured with HLA-A*11:01-expressing K562 cells pulsed with DMSO or AVEEVSLRK at 1 or 10 µg/mL. Heatmaps show IFN-γ production measured the following day using the U-PLEX assay. The mean of two technical replicates is shown. Panels A and B show results obtained using T cells from two independent healthy donors.

**Supplementary Figure 3.**
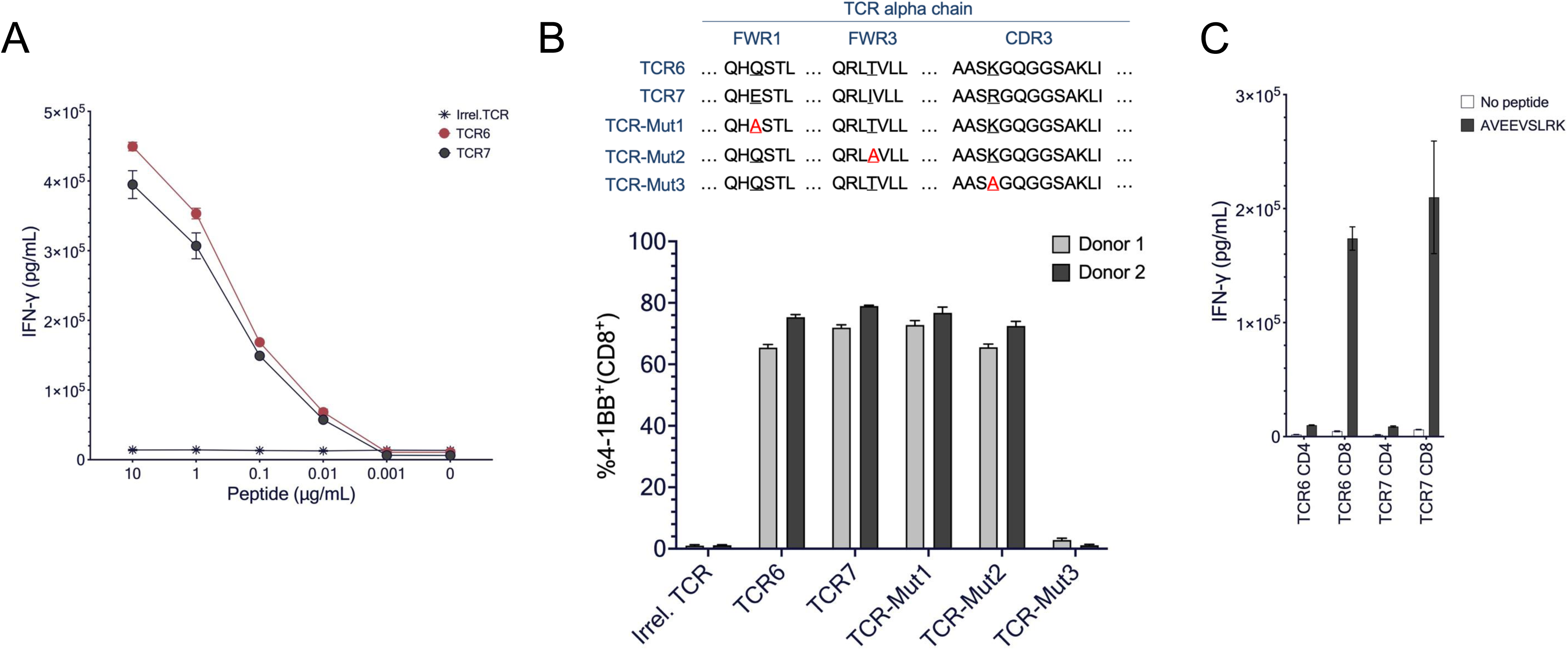
Additional data supporting Figure 4. **(A)** Human T cells from a donor different from that shown in Figure 4A were transduced with TCR6, TCR7, or an irrelevant TCR and co-cultured overnight with HLA-A*11:01-overexpressing K562 cells pulsed with decreasing concentrations of AVEEVSLRK. Line graph shows IFN-γ production measured using the IFN-γ U-PLEX assay. **(B)** To determine whether the three amino acid differences in the α chains of TCR6 and TCR7 had functional significance, three TCR6 variants were generated by substituting each differing residue with alanine (TCR-Mut1–TCR-Mut3). The first two residues were in framework regions 1 and 3 (FWR1 and FWR3), respectively, and the third in complementarity-determining region 3 (CDR3). The sequence differences and mutation strategy are shown in the upper panel. T cells from two healthy donors were transduced with an irrelevant TCR, TCR6, TCR7, or one of the three mutant receptors and co-cultured with K562-A*11:01 cells pulsed with AVEEVSLRK (0.1 µg/mL). Bar graphs show the percentage of TCR^+^ CD8^+^ T cells expressing the activation marker 4-1BB, as determined by flow cytometry the following day. **(C)** CD4^+^ TCR^+^ and CD8^+^ TCR^+^ cells isolated from TCR6-and TCR7-transduced human T cells from a donor different from that shown in Figure 4B were co-cultured overnight with K562-A*11:01 cells pulsed with AVEEVSLRK (1 µg/mL). Bar graph shows IFN-γ production measured using the U-PLEX assay. For all panels, the mean and standard deviation of two technical replicates are shown for each condition.

**Supplementary Figure 4.**
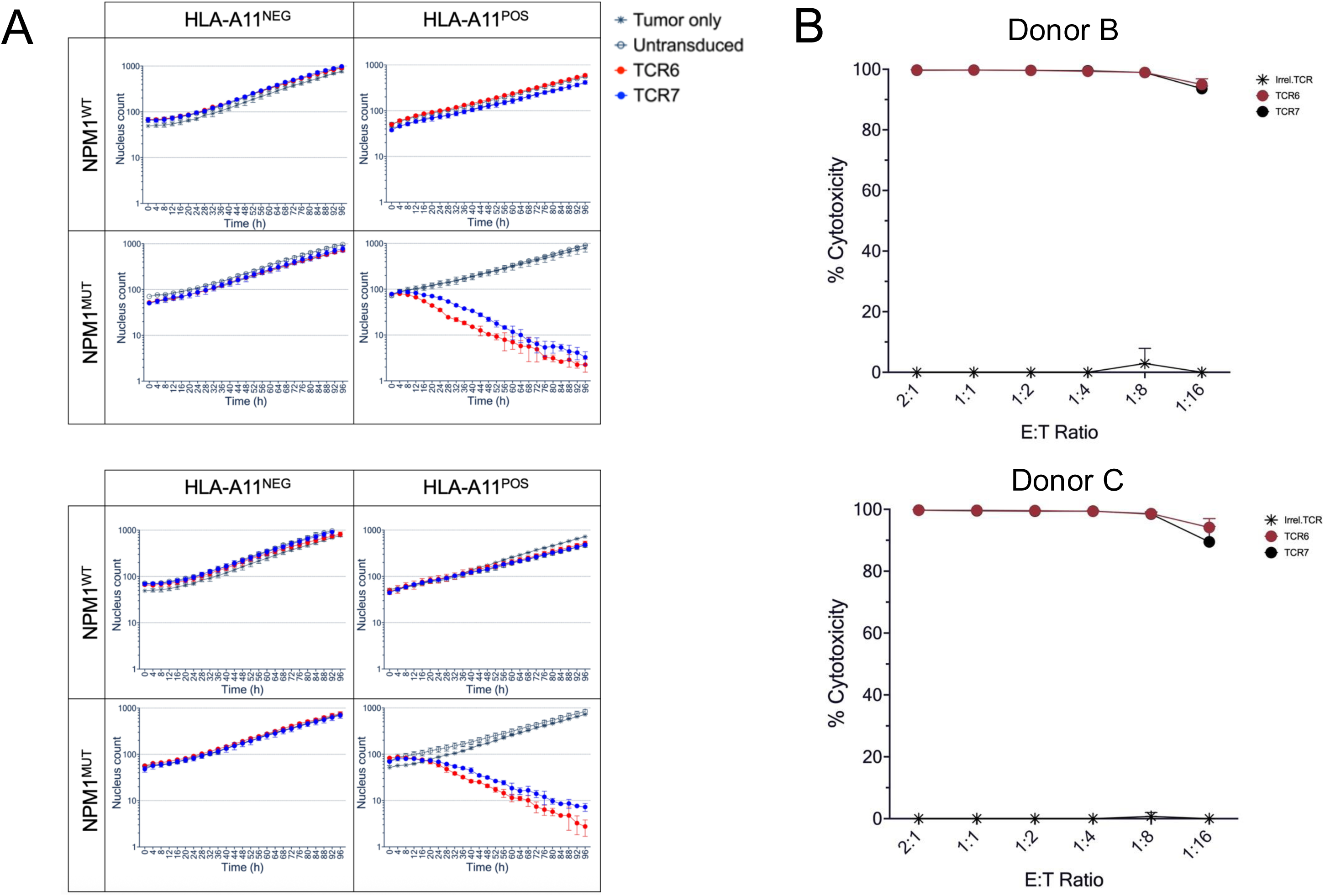
Additional data supporting the cell line testing shown in Figure 5. **(A)** Live-cell imaging data used to generate the cytotoxicity results shown in Figure 5A. Plots show the time course of co-cultures involving OCI-AML2 (NPM1^WT^) and OCI-AML3 (NPM1^MUT^) cells, with or without transduced HLA-A*11:01 (HLA-A11^POS^ or HLA-A11^NEG^). Two different healthy donors were used for this experiment (upper and lower panels). **(B)** Experiment performed as in Figure 5B using T cells from two different donors (upper and lower panels); these are representative experiments demonstrating activity at lower E:T ratios.

**Supplementary Figure 5.**
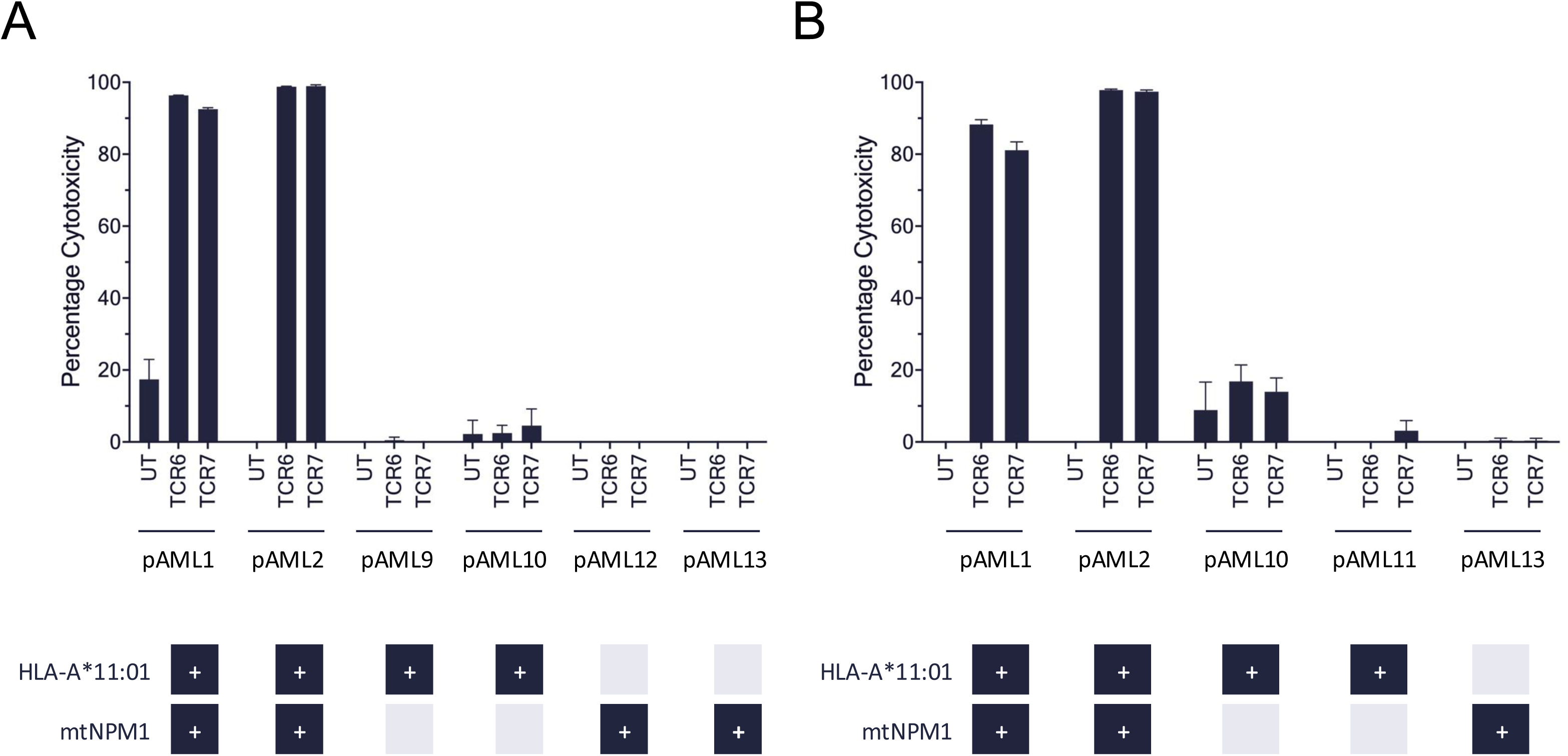
Additional data supporting the primary AML cell testing shown in Figure 5. Panels A and B show two additional independent experiments in which TCR6 and TCR7 were tested against primary AML samples as described in Figure 5C; a broader set of pAML samples was included in these experiments. The mean ± SD of three technical replicates is shown for each condition. Note that untransduced (UT) CD8^+^ T cells were used instead of irrelevant TCR-transduced cells for technical reasons.

**Supplementary Figure 6.**
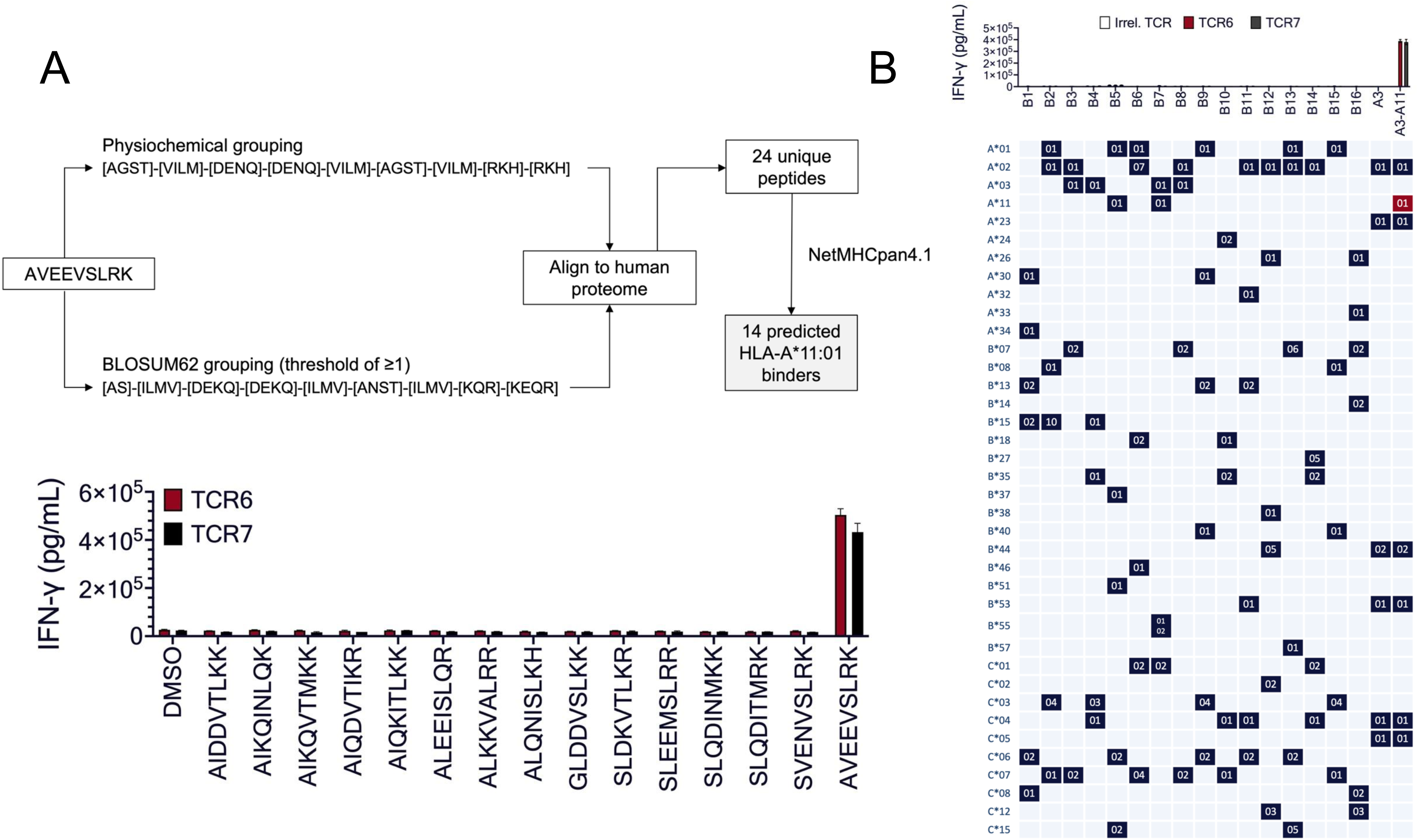
Additional cross-reactivity and alloreactivity data related to Figure 6. **(A)** To identify additional potentially cross-reactive peptides beyond those sharing the AVEEVSLRK recognition motif shown in Figure 6A, two strategies were used. First, an input sequence was derived from AVEEVSLRK based on physicochemical grouping, as described in reference 14. Second, a BLOSUM62-based strategy with a threshold of at least 1 was used to generate a second input sequence. These two inputs were then queried against ScanProsite, yielding 24 unique peptides in the human proteome, which were subsequently evaluated for HLA-A*11:01 binding using NetMHCpan-4.1. This identified 14 peptides with a binding rank below 2; these were considered binders. These peptides, together with AVEEVSLRK and DMSO, were pulsed onto K562-A*11:01 cells and co-cultured with TCR6- or TCR7-transduced T cells. Bar graph shows IFN-γ production measured by U-PLEX the following day. **(B)** TCR-transduced T cells were co-cultured with Epstein-Barr virus-transformed B-cell lines (EBV-B) derived from 16 HLA-typed healthy donors. HLA allele groups are shown in the leftmost column, and the specific HLA alleles for each donor are shown in the corresponding cells of the table below the graph. The graph above shows results of IFN-γ U-PLEX. The same controls as in Figures 6B and 6C were used. For both panels, mean ± SD of two technical replicates is shown. Results were validated using T cells from another healthy donor and in an independent experiment (not shown).

**Supplementary Figure 7.**
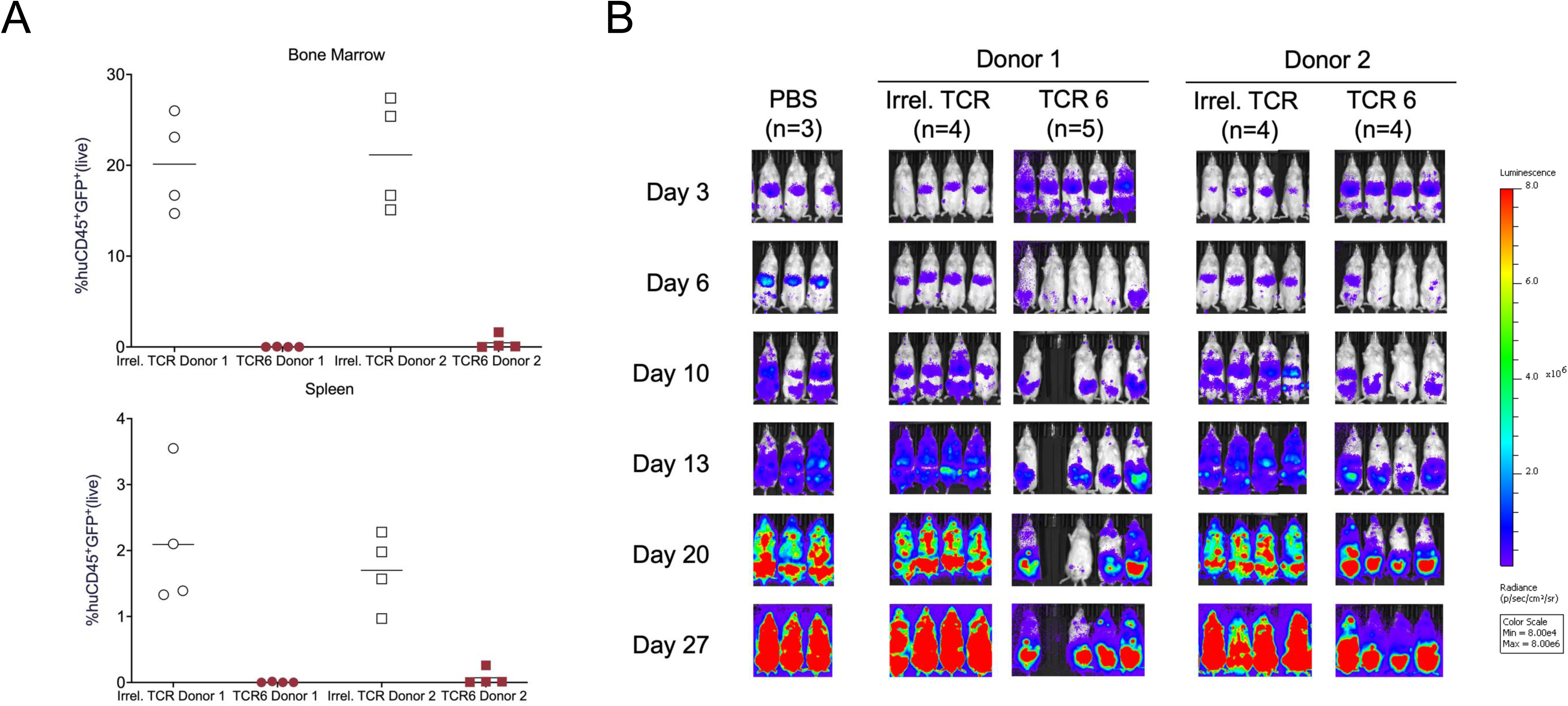
Additional adoptive cell transfer experiment. The experiment was performed as described in Figure 7; TCR-T cells were injected on day 3. To simplify the experiment, only TCR6 was tested, as TCR6 and TCR7 had shown similar performance in all prior experiments. **(A)** Flow cytometric analysis of bone marrow and spleen performed as described in Figure 7A. **(B)** Serial bioluminescence imaging.

**Supplementary Figure 8.**
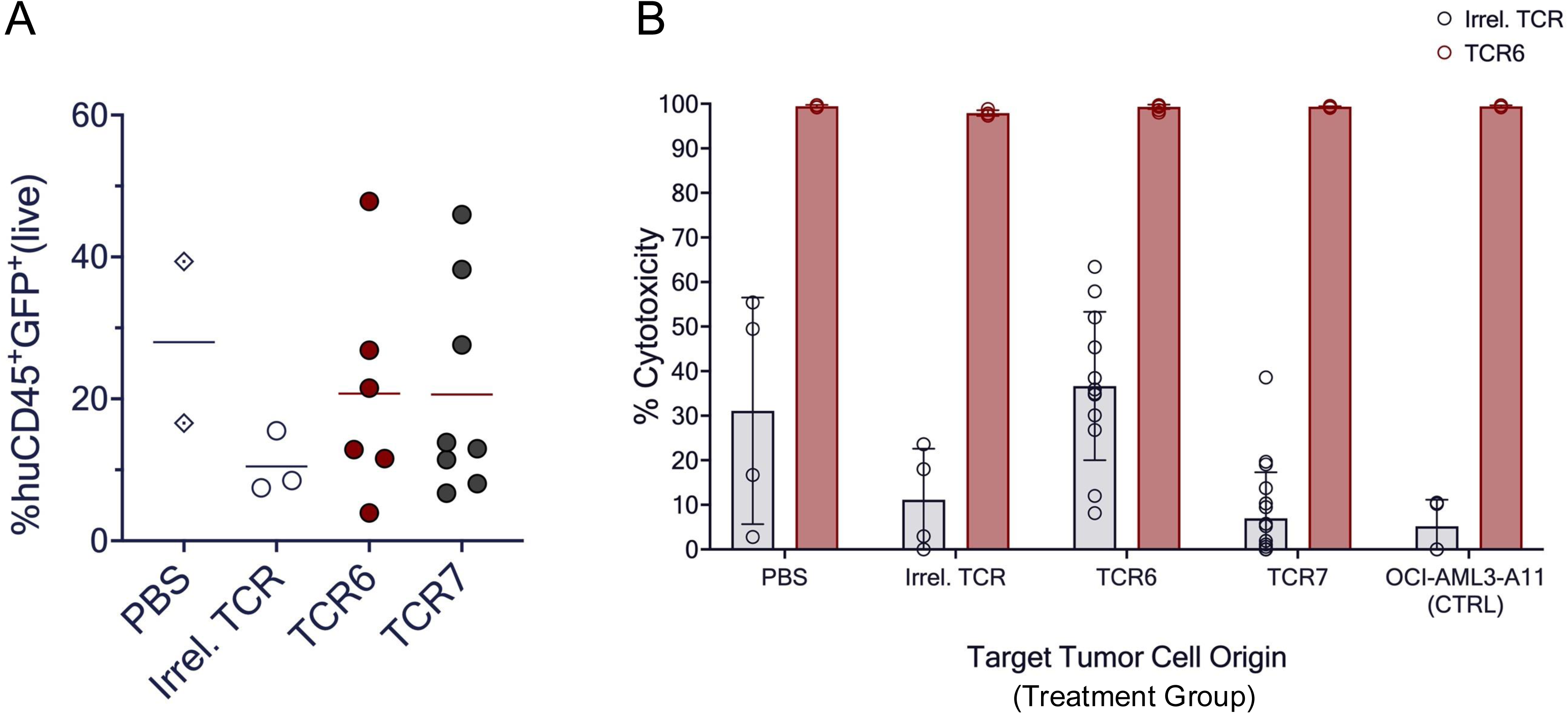
Additional data supporting Figure 7. **(A)** On day 35, mice described in Figure 7 were sacrificed. Large intraperitoneal tumors were noted in the pelvic region, although the exact organ of origin could not be determined. These were collected, homogenized, and analyzed by flow cytometry. Plots show the percentage of human CD45^+^ GFP^+^ AML cells among live cells for mice in each treatment group. Each symbol represents an individual mouse, and lines indicate mean values. For this analysis, data from the two donors were pooled; not all tumors were analyzed because of technical limitations. **(B)** Homogenized tumor cells from (A) were co-cultured with TCR6- or an irrelevant TCR-transduced T cells and quantified 72 hours later by flow cytometry based on human CD45 and GFP expression. Bar graphs show calculated percentage cytotoxicity for tumor cells from each treatment group (shown on the x-axis). GFP^+^ OCI-AML3-A*11:01 cells derived from the original cells used for mouse injection, but maintained *in vitro*, served as a positive control.

